# Heartbeat-evoked responses in M/EEG: A systematic review of methods with suggestions for analysis and reporting

**DOI:** 10.1101/2025.08.08.668923

**Authors:** Paul Steinfath, Maria Azanova, Nikolai Kapralov, Thomas Loesche, Lioba Enk, Vadim Nikulin, Arno Villringer

## Abstract

Heartbeat-evoked responses (HER), as measured by electroencephalography (EEG) or magnetoencephalography (MEG), represent neural activity time-locked to heartbeats and are widely used as a marker of cardiac interoception in the study of brain-body interactions. However, HER studies report largely variable findings, at least partially due to methodological variability. To achieve consensus on HER processing and improve the reproducibility of findings, the field urgently requires a structured summary of the methods employed so far. To this end, we conducted a systematic review of 132 HER studies using non-invasive M/EEG recordings in humans. Our results reveal substantial heterogeneity across most steps of HER analysis, ranging from data acquisition and preprocessing to HER estimation and statistical approaches. The large diversity in the processing choices is accompanied by considerable proportions of unreported methodological information across reviewed studies, reaching up to 80% for key processing steps. In addition, less than 33% of studies had enough statistical power to reliably detect meta-level HER effects, while their reported spatiotemporal locations varied substantially. We provide a comprehensive reporting and quality control checklist to aid in the development of more standardized procedures, highlighting critical steps for robust HER investigations. Additionally, we share the full extracted dataset, including an interactive version, to support other researchers in answering additional specific questions they may have. We hope that these resources will improve the robustness, reproducibility, and transparency of research in the growing HER field.

## 1. Introduction

The dynamic interplay between the central nervous system and other bodily organs is increasingly recognized as a crucial factor in understanding cognition, behavior, and overall well-being (Azzalini et al., 2019; Criscuolo et al., 2022; Villringer et al., 2025). A growing body of research has demonstrated how the interaction between the brain and body shapes many functional aspects, ranging from the direct modulation of neuronal activity by blood pressure (Jammal Salameh et al., 2024) up to higher cognition (Umeda et al., 2016) and complex emotional states (Feldman et al., 2024; Garfinkel et al., 2014).

Among the various organs involved, the heart plays a particularly important role in brain-body interaction research. Beyond its essential function of delivering oxygenated blood and vital nutrients to the brain and the rest of the body (Hall & Hall, 2021), its rhythmic activity itself appears to serve as a pacemaker, providing an important reference frame for the brain (Tallon-Baudry et al., 2018). Accumulating evidence suggests that distinct phases of the cardiac cycle are advantageous for either perception or action; therefore, monitoring and adjustment of cardiac activity by the brain might be necessary to achieve optimal performance (Galvez-Pol et al., 2022; Skora et al., 2022).

One key methodology for studying heart-brain interactions with electroencephalography (EEG) or magnetoencephalography (MEG) involves analyzing neural activity time-locked to the heartbeat. The resulting heartbeat-evoked responses^1^ (HERs) reflect the processing of heartbeat-related information in the central nervous system (Dirlich et al., 1997; Schandry et al., 1986). HERs are commonly used as a marker for cardiac interoception (Pollatos & Schandry, 2004) and have been linked to a broad range of functions, such as perception (Al et al., 2020; Park et al., 2014), attention (Banellis & Cruse, 2020), motor activity (Al et al., 2023), or emotions (Di Bernardi Luft & Bhattacharya, 2015).

Despite the growing interest and number of studies, the field of HER research is facing a challenge of methodological heterogeneity and variability in reported findings (Coll et al., 2021). There appears to be no single, unified manifestation of HER. Instead, heartbeat-locked neural activity has been associated with a wide range of cognitive and emotional functions and observed across diverse topographic and temporal regions (Engelen, Solcà, et al., 2023; Park & Blanke, 2019). While some of this variability can be attributed to participant characteristics (e.g., patients or controls) or the experimental context in which HERs are investigated, variations in data acquisition, preprocessing techniques, and statistical analyses likely also play a role.

Beyond the general methodological decisions that apply to all studies of evoked responses (such as how to filter the data and whether to use baseline correction), analysis of HERs requires special treatment, most notably due to the cardiac field artifact (CFA). CFA is caused by the instantaneous spread of magnetic and electric fields from the heart’s activity due to volume conduction, and is directly captured by M/EEG sensors (Dirlich et al., 1997). In classical task-based studies, the CFA is widely believed to play a minor role, as cardiac-related noise is not time-locked to stimuli and thus suppressed by trial averaging. However, due to the inherent coupling between heartbeat and CFA, it is not reduced by averaging across trials when obtaining HERs and poses a serious challenge for HER research. To investigate the brain activity related to the heartbeat, without the CFA, researchers typically either try to avoid time ranges corresponding to high CFA, remove the CFA from the data, or include the heart’s activity as a confound in the statistical analysis. Various approaches are employed for this purpose, and the outcome may depend on the selected approach and even on decisions such as the choice of settings for electrocardiogram (ECG) recording (Gray et al., 2007). To note, other confounds, such as pulse-related movement of tissue and sensors, can also introduce heartbeat-locked artifacts (Kern et al., 2013), but their influence on measured HERs remains unclear.

Apart from cardiac artifacts, the overlap of heartbeat- and task-evoked activity additionally complicates the analysis of HERs. When HERs are studied in task paradigms, the potential overlap with task-evoked activity must be accounted for (Azzalini et al., 2019; Steinfath et al., 2025). Furthermore, due to the cyclic nature of the heartbeat, neural responses to successive heartbeats can superimpose, especially when inter-beat intervals (IBIs) are short. The necessity to address these problems further increases the number of analysis decisions researchers have to make.

Overall, it remains unclear to what extent methodological choices influence the HER effects reported across different studies, and thorough examination of the employed methods has been repeatedly advocated (Coll et al., 2021; Park & Blanke, 2019). This situation mirrors the reproducibility challenge in the broader field of neuroscience (Botvinik-Nezer & Wager, 2023), which is currently emphasized and addressed by newly appearing analytic approaches such as large-scale collaborative data analyses (EEGmanypipelines, Trübutschek et al., 2024), multiverse analyses (Clayson et al., 2021), or meta-methods reviews (Elson, 2019).

In light of these challenges, it is an essential first step to systematically collect and assess the methodological approaches used in HER research. To achieve improved consistency and reproducibility in HER analysis, a clear understanding of current practices is necessary. This review, therefore, focuses specifically on the methods used in HER research. To this end, we extracted and analyzed the methodological decisions, from data acquisition through preprocessing to statistical analysis, from published HER studies.

In addition, we highlight aspects that may be especially important for the HER analysis, which we hope will stimulate further HER research, discussion, increased transparency, and potential consensus. To facilitate this, we publicly share the full resulting dataset, its interactive version, the accompanying codebook, and the HER analysis checklist with this article.

## 2. Methods

### 2.1. Literature search

We conducted a systematic literature search using the PubMed database on three separate dates to identify studies investigating HERs using noninvasive neuroimaging. Table 1 specifies the search terms used, with corresponding dates and the number of studies retrieved, as well as the total number of studies after duplicate removal.

**Table 1.** Search strategy.

| Date | Search query | Studies retrieved |
| --- | --- | --- |
| 15.01.2024 | (eeg OR electroencephalograph*) AND<br>(heartbeat evoked potential OR heartbeat evoked response)<br>NOT (Review[Publication Type] OR Systematic<br>Review[Publication Type]) AND English[Language] | 647 |
| 24.07.2024<br>(rerun:<br>05.08.2024) | (eeg OR electroencephalograph* OR meg OR<br>magnetoencephalograph*) AND (heartbeat evoked potential<br>OR heartbeat evoked response) NOT (Review[Publication<br>Type] OR Systematic Review[Publication Type]) AND<br>English[Language] | 682<br>(rerun:<br>683) |
| 05.08.2024 | ("heartbeat evoked" OR "heart evoked") AND Journal<br>Article[Publication Type] AND English[Language] | 166 |
| **Total amount of unique studies after removing duplicate entries** |  | **749** |

After the removal of duplicates, all retrieved records (N = 749) were screened following the guidelines of the Preferred Reporting Items for Systematic Reviews and Meta-Analyses (PRISMA; Moher et al., 2009). We included studies that: (1) examined HERs, (2) applied noninvasive EEG or MEG recordings, (3) were conducted in human participants, (4) reported original data, (5) were in English language, and (6) had full text available. Exclusion criteria were: (1) intracranial recordings (due to the different data analysis approaches), (2) non-original research (reviews, commentary papers, and toolboxes), (3) studies not analyzing evoked responses (e.g., time-frequency analysis only), and (4) lack of a peer review. Because this review focuses on HER analysis methods, we also included studies analyzing data in source space and those without statistical testing. After screening titles and abstracts, we evaluated the full texts of potentially relevant articles to determine their eligibility. After applying the above-mentioned criteria, 127 articles were included. In addition, we identified seven references through the bibliographies of the reviewed publications, of which five were included, resulting in a final sample of 132 articles used in the systematic review. The screening process is detailed in the PRISMA flow diagram (Fig. 1A).

**Figure 1.**
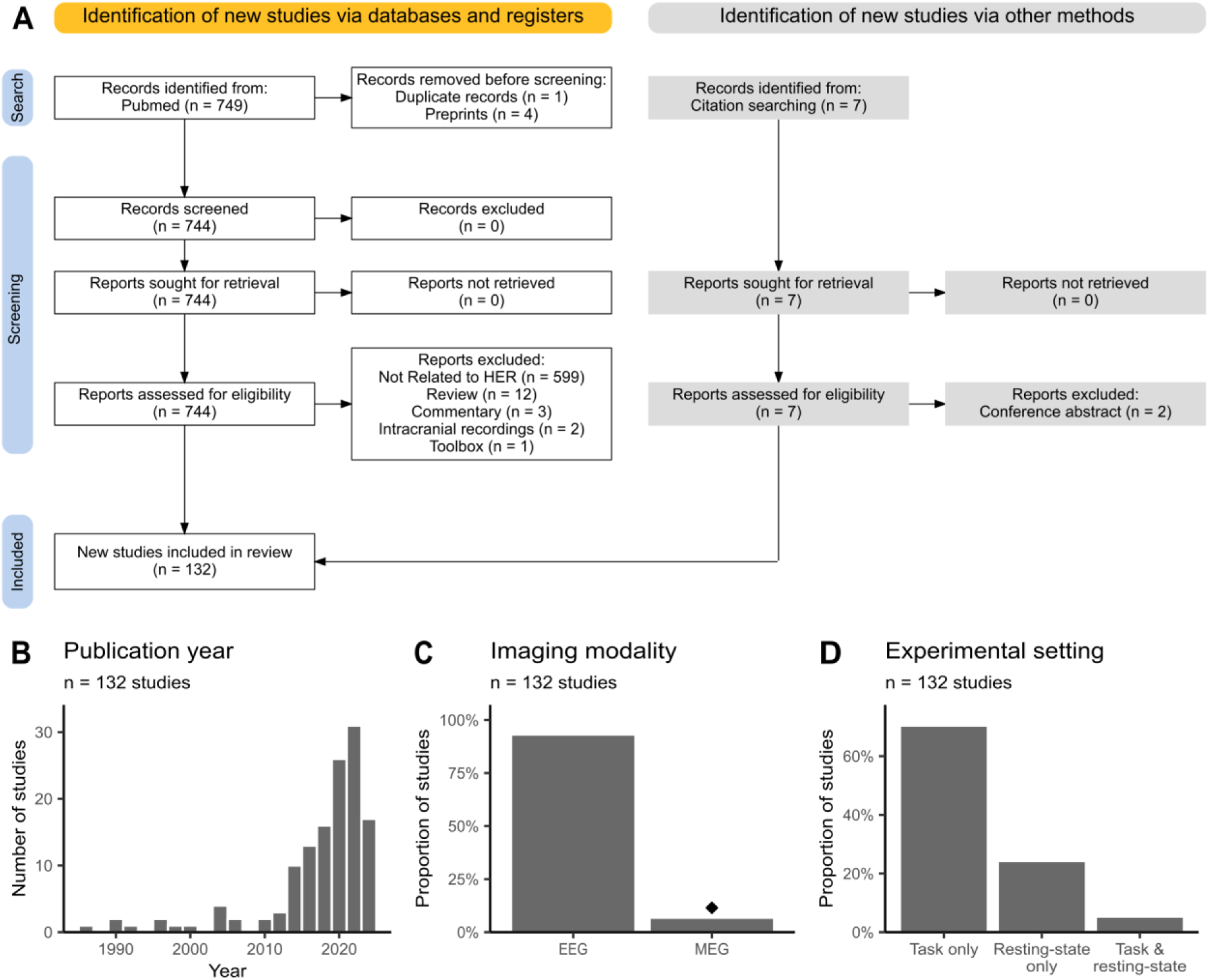
Overview of the studies included in the review. (A) Flowchart describing the literature selection process according to the PRISMA guidelines. (B) The majority of the included studies were published within the last 10 years. (C) Most studies use EEG data to analyze heartbeat-evoked responses (HERs). Diamond indicates that most MEG studies come from one research group. (D) HERs are typically analyzed during task performance rather than in the resting state.

### 2.2. Extraction of methodological choices

In the following, we use the term *processing steps* when referring to the various data transformations or parameters that researchers choose (for example, the cutoff frequency of a high-pass filter). A particular value (e.g., 1 Hz) that is then selected for a processing step is referred to as a *(methodological) choice*, and a combination of methodological choices for all processing steps is considered as a *pipeline* in this review. If a study uses different methodological choices for any processing step (e.g., performs the analysis with and without baseline correction), we say that several pipelines correspond to this same study.

For each article included in the review, we extracted methodological choices for a wide range of processing steps, from the acquisition of physiological data (EEG/MEG/ECG) to the calculation and statistical analysis of HERs (Table 2). Three reviewers, independently, extracted the data and then additionally checked for the consistency of the extracted information for each processing step. We developed a standardized codebook to unify and ensure consistency of labels referring to the same methodological categories (e.g., “linked mastoids” and “averaged mastoids”). The codebook is publicly available and can be found together with the final dataset at https://osf.io/znrbm.

**Table 2.**
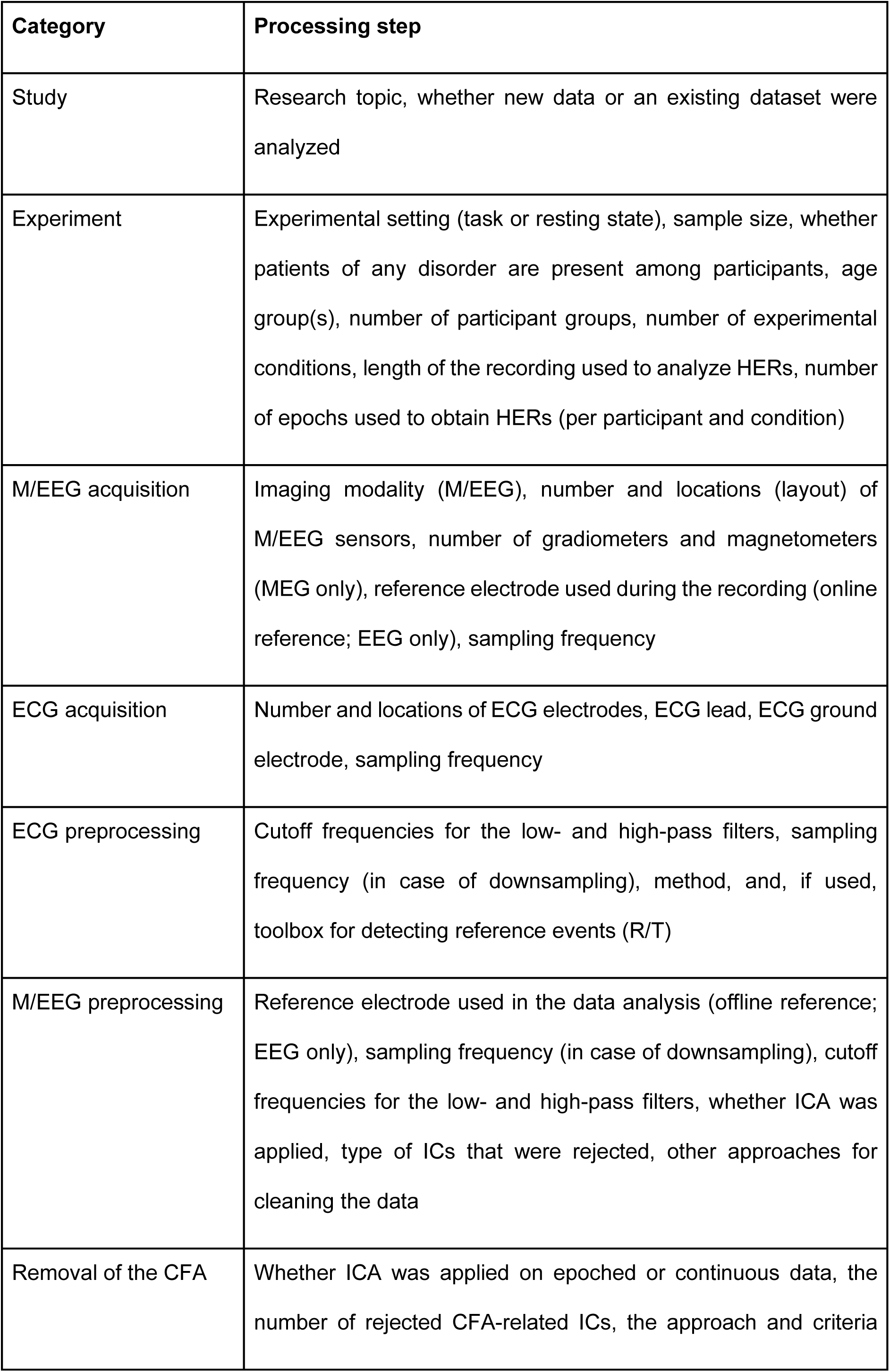
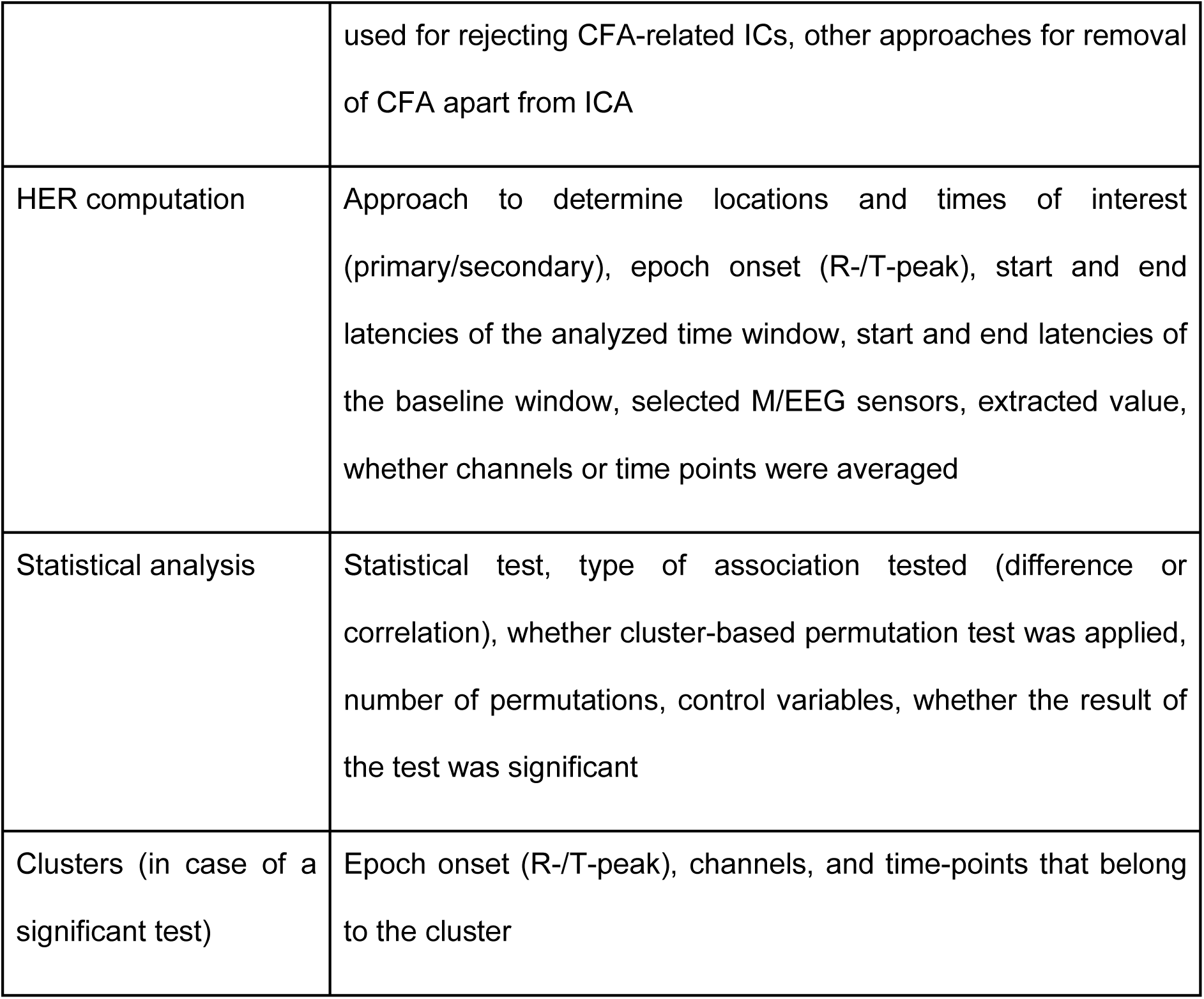
Overview of the extracted processing steps from pipelines used for studying HERs. Abbreviations: HERs – heartbeat-evoked responses, M/EEG – magneto- or electroencephalography, ECG – electrocardiogram, ICA – independent component analysis, ICs – independent components, CFA – cardiac field artifact.

| Category | Processing step |
| --- | --- |
| Study | Research topic, whether new data or an existing dataset were analyzed |
| Experiment | Experimental setting (task or resting state), sample size, whether patients of any disorder are present among participants, age group(s), number of participant groups, number of experimental conditions, length of the recording used to analyze HERs, number of epochs used to obtain HERs (per participant and condition) |
| M/EEG acquisition | Imaging modality (M/EEG), number and locations (layout) of M/EEG sensors, number of gradiometers and magnetometers (MEG only), reference electrode used during the recording (online reference; EEG only), sampling frequency |
| ECG acquisition | Number and locations of ECG electrodes, ECG lead, ECG ground electrode, sampling frequency |
| ECG preprocessing | Cutoff frequencies for the low- and high-pass filters, sampling frequency (in case of downsampling), method, and, if used, toolbox for detecting reference events (R/T) |
| M/EEG preprocessing | Reference electrode used in the data analysis (offline reference; EEG only), sampling frequency (in case of downsampling), cutoff frequencies for the low- and high-pass filters, whether ICA was applied, type of ICs that were rejected, other approaches for cleaning the data |
| Removal of the CFA | Whether ICA was applied on epoched or continuous data, the number of rejected CFA-related ICs, the approach and criteria |
|  | used for rejecting CFA-related ICs, other approaches for removal of CFA apart from ICA |
| HER computation | Approach to determine locations and times of interest (primary/secondary), epoch onset (R-/T-peak), start and end latencies of the analyzed time window, start and end latencies of the baseline window, selected M/EEG sensors, extracted value, whether channels or time points were averaged |
| Statistical analysis | Statistical test, type of association tested (difference or correlation), whether cluster-based permutation test was applied, number of permutations, control variables, whether the result of the test was significant |
| Clusters (in case of a significant test) | Epoch onset (R-/T-peak), channels, and time-points that belong to the cluster |

### 2.3. Data analysis

Data analysis and visualization were performed using R (version 4.4.3; R Core Team, 2025). For most processing steps, we assessed the overall scope of methodological choices and the proportion of studies or pipelines that utilized each choice. Since the information relevant for this review is not reported for some studies and pipelines, we specify the exact number of data points above each plot. Some studies perform several analyses with different selections of methods — if this is the case for a particular plot, we refer to the data points as pipelines; otherwise, we refer to them as studies. Importantly, the number of studies/pipelines using a specific methodological choice can increase not only because of community consensus, but also when the same research group repeats acquisition or analysis settings across multiple studies. To address it in histogram plots, we add a diamond symbol above choices, for which the same first or last author contributes at least 50% of the occurrences.

#### Entropy

To quantify and compare the variability of methodological choices for different processing steps, we used the concept of Shannon entropy (Shannon, 1948). Let *k* denote the number of possible choices, while p_1,_ p_2_,…, p_k_ stand for their occurrence proportions across studies or pipelines. Then, Shannon entropy is computed as:

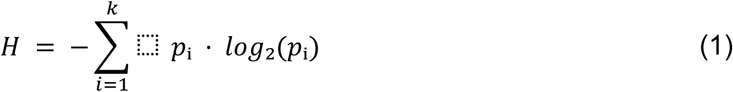

Larger entropy values indicate a greater heterogeneity in the choices, while entropy is zero when all studies use the same method. As the maximum possible entropy depends on the number of available options, we normalized it to allow for comparisons across processing steps with different numbers of choices:

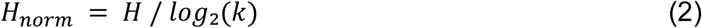

Additionally, we binned continuous parameters (e.g., the number of trials) that contained more than 10 unique values to make them comparable to categorical variables before computing entropy.

#### Aggregation of EEG locations

To analyze the spatial distribution of observed HER effects, we collected information about the EEG channels included in the HER analysis and, if applicable, about those belonging to at least one of the clusters reported for significant tests. To combine studies that used different EEG layouts during data recording, we calculated the number of occurrences for each channel and layout separately. The occurrence counts were interpolated to form topographies independent of actual sensor locations. The topographies were then summed across the considered layouts, that is, variations of the 10-20 system with 19, 32, and 64 channels, as well as the 128-channel BioSemi layout.

#### HER peak timing comparison

To statistically compare the timing of effects observed with different HER analysis strategies (e.g., clustering vs. averaging), we implemented a permutation test. For each strategy, the distribution of analysis time window start and end times was aggregated as a cumulative sum, and the peak was identified. Next, the peak time difference between the two strategies was calculated. To test if the differences between strategies were statistically significant, all analysis time windows from both strategies were pooled and randomly assigned to two groups of the original sizes. The peak timing differences were recalculated, and the permutation procedure was repeated 1000 times. Lastly, p-values were obtained by comparing the original peak timing difference to the permutation distribution.

#### Definition of cluster-based tests

Note that throughout the manuscript, by “cluster-based permutation tests” (also “permutation clustering”, “cluster-based analysis”, and similar) we specifically refer to cluster-sum tests, which establish significance based on the summed statistic (e.g., t-values) across spatiotemporal points within a cluster (Maris & Oostenveld, 2007). While this approach is currently popular in the M/EEG field, it comes with the limitation that the precise timing and spatial extent of clusters should be interpreted with caution (Sassenhagen & Draschkow, 2019). Other cluster-based approaches exist (Frossard & Renaud, 2022; Rousselet, 2025), but, due to their novelty and lack of implementation in neuroscientific toolboxes, their prevalence in HER research was not assessed in this review.

#### Minimal detectable effect sizes

To compare minimal effect sizes that could be robustly detected across pipelines, we estimated the smallest effect that a study can reliably identify given a specified level of confidence. Such estimates provide a practical way to assess whether a study is sufficiently powered for the effects it aims to detect. Accordingly, we (1) computed minimal detectable effect size for one- and two-sample t-tests, ANOVAs, and correlations using pwr R package (version 1.3-0; Champely et al., 2020) based on the information about sample sizes, numbers of groups, conditions, and required power of 80% with significance level set at 5%, (2) converted these effect sizes to Cohen’s d, when required, and finally (3) converted Cohen’s d into Hedges’ g using the exact correction approach (Hedges & Olkin, 1985). Hedges’ g was used instead of Cohen’s d because it provides a less biased effect size estimate for small sample sizes, which are typical for HER studies. The resulting Hedges’ g estimates of minimal detectable effect size were compared to previously reported effect sizes in HER literature (Coll et al., 2021). Details on the estimation of Hedges’ g are provided in Section A of the Supplementary Material. Finally, we examined how detectable effect sizes could change if within-participant variability in HER values across epochs was taken into account (see Section B of the Supplementary Material for more details on these estimations).

#### Estimation of the number of epochs

As M/EEG signals are prone to high amounts of environmental and physiological noise, obtaining a clear evoked response often requires averaging over a substantial number of epochs. For each pipeline, we extracted the minimum (preferred) or mean number of heartbeat-locked epochs per participant and condition, where possible. Those were not readily available in some studies. For such cases, we estimated approximate numbers using strategies such as (1) dividing the overall reported number of epochs by the number of analyzed participants, (2) multiplying the reported length of the recording in minutes within which HERs were analyzed by the average heart rate, when reported, or otherwise by 60 beats per minute. If reported, those were adjusted by the number of epochs removed during cleaning.

## 3. Results

The results of the review are structured as follows. We start with an overview of included studies (Section 3.1), followed by an overview of processing steps that were extracted and analyzed (Section 3.2). Sections 3.3-3.9 provide a detailed description of methodological choices that occur in the literature for various processing steps from data acquisition to statistical analysis.

### 3.1. Overview of the included studies

In total, 132 studies published between 1986 and 2024 were included in the review (Fig. 1B). The vast majority of studies used EEG recordings (123/132, 93%), while only nine studies (6.8%) employed MEG to measure HERs (Fig. 1C). As shown in Fig. 1D, most studies (93/132; 70%) focused on the analysis of HERs during performance of various tasks involving perception of own heartbeats (see Coll et al., 2021 for a review) or external stimuli (e.g., Al et al., 2020; Marshall et al., 2020; Park et al., 2014). A smaller proportion of studies (32/132; 24%) performed resting-state analyses, testing the potential relationship between the amplitude of HER and diverse psycho-physiological states (e.g., Flasbeck et al., 2021; Giusti et al., 2024; Yoris et al., 2024) as well as diverse clinical conditions (e.g., Birba et al., 2022; Kumral et al., 2022; Schmitz et al., 2021; Solcà et al., 2020). Seven studies (5.3%) analyzed HERs both during task and resting state.

The included studies were published in a wide range of journals (Table S2) and varied in the patient or age groups studied, research topics, imaging modalities, and specific HER research groups. All these factors likely contribute to the variability in methods and findings that we document in this review and may affect some processing choices more than others. We invite readers to use the provided interactive resource (https://paulsteinfath.shinyapps.io/her-systematic-review/) to investigate the importance of these factors, which are outside the scope of the current review.

### 3.2. Overview of methodological choices

For each processing step of Table 2, we extracted methodological choices made by researchers (e.g., which value of a parameter to use or whether to apply a certain method). In total, we extracted 410 pipelines (note that some studies included multiple pipelines due to the use of different parameters or strategies). In the subsequent sections, we only consider distinct pipelines with respect to each processing step we discuss. For example, when discussing preprocessing steps, multiple pipelines that differ only in terms of statistical analysis are treated as one. This should ensure that our analyses reflect methodological variability within each processing step without being inflated by differences in unrelated steps. The total number of pipelines (or studies, when only one pipeline per study was reported) is specified above each figure panel or in parentheses after each reported value. Note that several resulting pipelines may come from one laboratory (max. 12 studies from the same last author) or from re-analyses of the same dataset (max. 2 studies).

Based on the information we extracted for data processing steps, from acquisition to statistics, we observe a substantial lack of consensus in most methodological choices. We quantify this heterogeneity for each processing step using normalized Shannon entropy (Eq. 2), where values close to 1 indicate high variability in choices, and values closer to 0 indicate a single dominant choice. Most steps showed large entropy (Fig. 2A). For example, frequency cutoffs of high- and low-pass filters used in preprocessing show great variability (Fig. 3E). Similarly, researchers select widely varying time windows of interest for their HER analysis (see Section 3.7 for more details). Few steps show consistency across the field: most studies use EEG data, select the R-peak as the reference event for time-locking HERs, and analyze HER amplitude.

**Figure 2.**
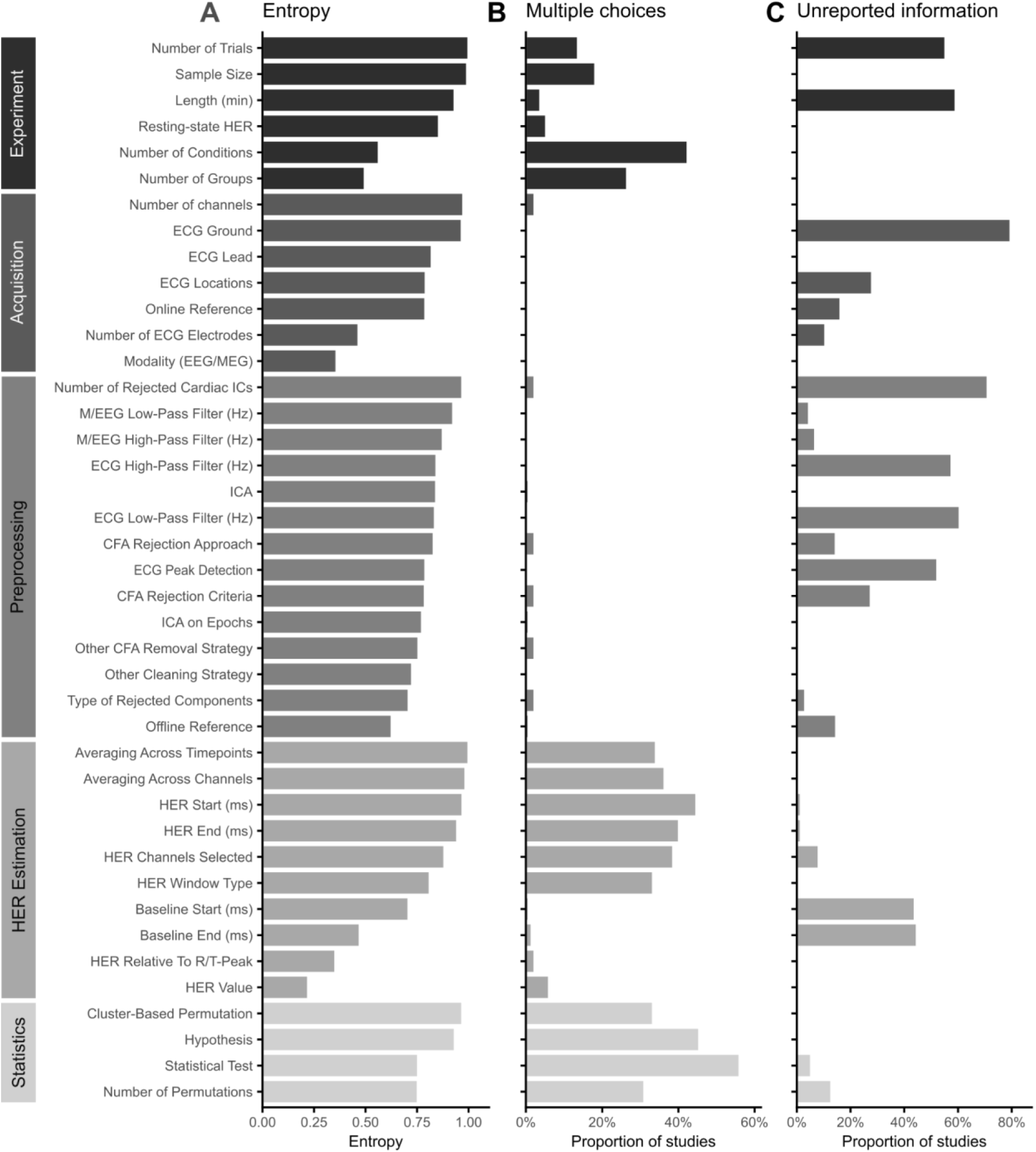
Overview of the extracted processing steps, heterogeneity in methodological choices, and unreported information. (A) Entropy was calculated to illustrate the diversity in processing choices across studies. Within categories, the processing steps are sorted by entropy in descending order. (B) Some studies use several analysis approaches, specifically to estimate heartbeat-evoked response (HER) values and perform the statistics. (C) The highest percentage of unreported information was observed for ECG ground location and the number of rejected cardiac independent components (ICs), a relevant parameter for artifact correction in the HER field. Abbreviations: ICA – independent component analysis, CFA – cardiac field artifact, ECG – electrocardiography, EEG – electroencephalography, MEG – magnetoencephalography.

**Figure 3.**
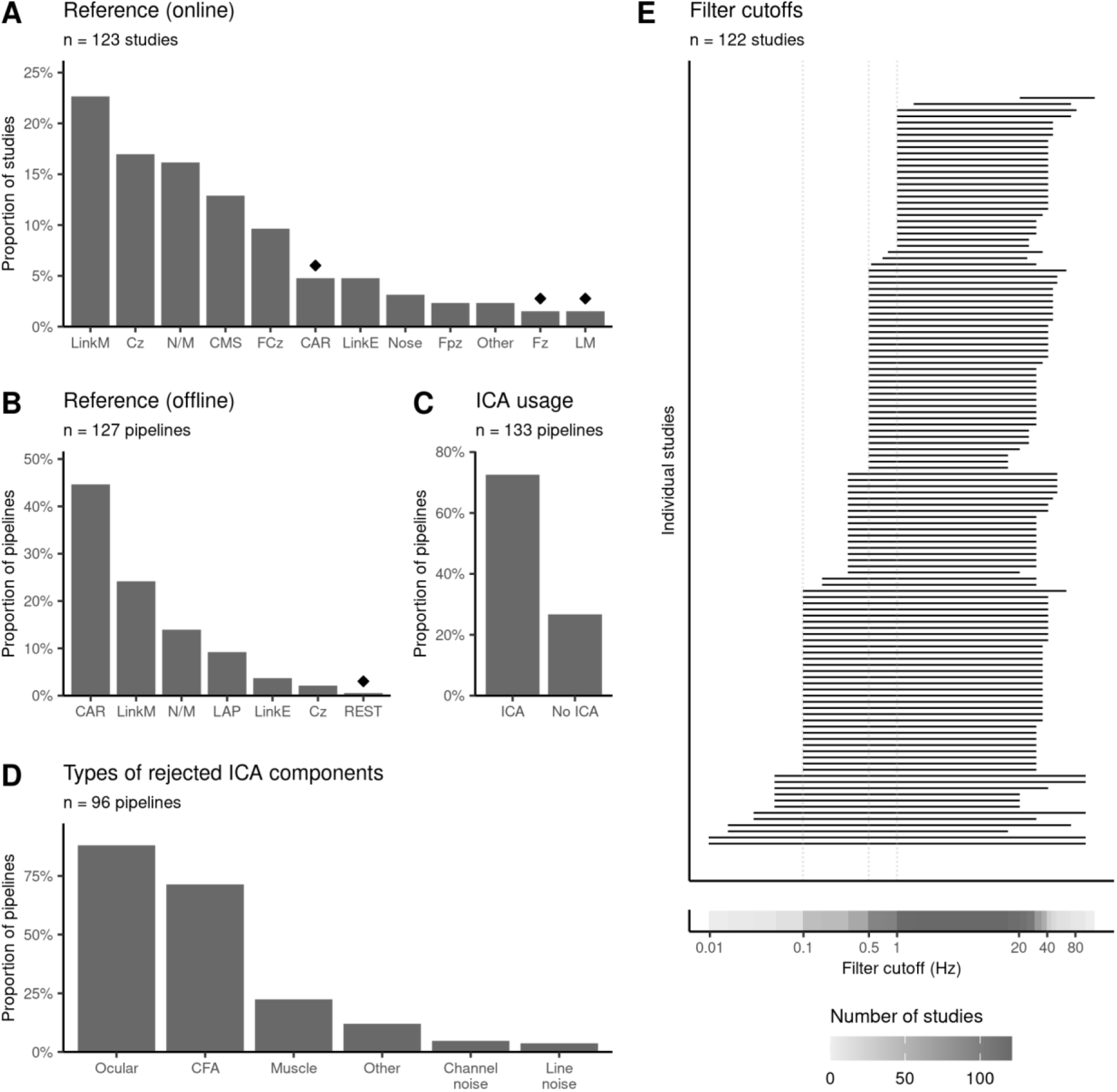
Heterogeneity in M/EEG acquisition settings and preprocessing methods. (A) Linked mastoids and Cz are most commonly used during the recording as reference. Only studies that used EEG are shown. The category Other denotes references that appeared only in a single study. Diamond symbols indicate choices driven by one research group (at least 50% of occurrences). (B) Common average is the most common reference used in the data analysis. Only studies that used EEG are shown. (C) Most studies use ICA for artifact removal. (D) ICA components related to eye movements (ocular) and CFA are rejected most frequently. (E) Frequency ranges analyzed by the reviewed studies. Popular high-pass filter choices include 0.1, 0.5, or 1 Hz. Abbreviations: LinkM – linked mastoids, CMS – common mode sense, CAR – common average reference, LinkE – linked earlobes, LM – left mastoid, LAP – laplacian or current source density analysis, N/M – not mentioned, REST – reference electrode standardization technique, ICA – independent component analysis, CFA – cardiac field artifact.

During acquisition and preprocessing, most studies apply exactly one choice per step, e.g., they either perform ICA for artifact correction or do not, but rarely both (Fig. 2B). In contrast, using several approaches for HER computation and statistical analysis within the same study is quite common: 56% of studies (74/132) use multiple statistical tests, and 45% (59/132) vary the start or the end latency of the time window of interest. Many such cases involve primary and secondary data analysis, or several related hypotheses within the same study, and therefore, typically do not require additional statistical correction.

However, almost no study systematically compares several methodological choices per processing step to assess the effects of researcher-dependent decisions on HER results. Among the reviewed studies, Candia-Rivera, Catrambone, et al. (2021) is a single such study that varies a preprocessing step (offline reference). Future studies should address the outstanding question of how processing decisions affect HER findings. Additionally, HER research articles could show if findings are robust to decisions such as baseline correction or different CFA cleaning methods.

Adding to that, a substantial proportion of methodological decisions is unreported (Fig. 2C). In 71% of studies reporting CFA-related independent component rejection (49/69), information on the number of rejected components is not provided. In 80% of studies (105/132), the ECG ground location is not explicitly stated. As natural contamination with CFA makes HERs distinct from other event-related responses, it is critical to provide as much information as possible about heartbeat-related processing and acquisition decisions. In addition, we observe a high percentage of unreported values for the following critical methodological choices: the number of epochs averaged to obtain HERs (73/132 studies, 55%), recording length (78/132, 59%), cutoff frequencies of high-pass (76/132, 58%) and low-pass (80/132, 61%) filters applied to ECG, algorithm for detection of R-/T-peaks in the ECG (69/132, 52%), and the start or end of the baseline window for HER (59/132, 45%). In 28% of studies (37/132), ECG locations are not reported. Overall, these results highlight that even for parameters critical to HER computation and analysis, there is a lack of transparency in the field, which contributes to reduced reproducibility of results. To remedy this issue, we provide a comprehensive reporting and quality control checklist with this review (Box 1).

### 3.3. M/EEG acquisition and preprocessing of evoked responses

Since HERs typically occur in the 0.5–2 μV range (Pérez et al., 2005; Pollatos & Schandry, 2004; Schandry & Weitkunat, 1990), M/EEG recordings of these signals often have a relatively low signal-to-noise ratio (SNR), requiring careful data preprocessing and artifact removal strategies. At the same time, the preprocessing itself can affect the shape and amplitude of ERP components (Clayson et al., 2021). In this section, we focus on the processing steps that are common to all ERPs, while the specific methods for suppressing CFA are discussed in more detail in Section 3.5.

In EEG, the choice of the reference electrode is important as it affects the shape and distribution of the ERP (Luck, 2014), making direct comparisons across studies with different references difficult. As shown in Fig. 3A, we observe some variability with respect to the online reference used during data acquisition, with linked mastoids (23%, 28/123 EEG studies) and Cz (17%, 21/123) as the most common options. This variability is reduced during data processing, where the common average reference (CAR, Fig. 3B) emerges as the most dominant choice (45%, 57/127 pipelines). This observation is complemented by Candia-Rivera, Catrambone, et al. (2021), who compared the influence of different re-referencing schemes on HERs and concluded that CAR leads to the most consistent results across a range of brain-heart interaction measures. However, they also note that the optimal re-referencing choice depends on the experimental settings and can vary across studies.

When analyzing relatively slow evoked responses, such as the HER, optimal filter settings are an important consideration to avoid removing or inducing effects (Luck, 2014; Tanner et al., 2015). We observe that, across studies, filtering is generally done between 0.1 and 40 Hz, with the most dominant high-pass filter cutoffs at 0.1, 0.5, and 1 Hz (Fig. 3E). In standard ERP analyses, frequencies above 40 Hz are typically excluded because they often are contaminated with different artifacts like muscle or line noise (Luck, 2014). Optionally, the data can be downsampled, but the final sampling frequency should be at least twice as high as the maximal frequency present in the recordings to avoid aliasing (e.g., at least 80 Hz for a 40 Hz low-pass filter). As shown in Fig. S3, all reviewed studies used sufficiently high sampling frequencies (at least 100 Hz) during recording and in offline analysis, compared to the popular choice of a 40 Hz low-pass filter.

Meanwhile, the high-pass filter cutoff must balance the removal of low-frequency noise while preserving the slowly evolving components of the heartbeat-locked signal. Given that some neurons track changes in inter-beat intervals (De Falco et al., 2024) and assuming the resting heart rate in humans is approximately 1 Hz, a high-pass filter of 1 Hz might inadvertently remove genuine HER activity. Ultimately, the filter selection represents a trade-off between the specific aims of the HER analysis and the noise characteristics of the dataset. Future studies are needed to shed light on the frequency components of the HER across contexts, their relationship to heart rate, and the impact of different filter cutoffs on the observed associations between HERs and behavior or cardiac artifacts. Approaches such as multiverse analysis could be used for the systematic exploration of such effects (Clayson et al., 2021). In the meantime, we recommend reporting the filter settings used to obtain HER (Box 1).

Due to the low amplitude of HERs, additional artifact-reduction strategies are often used to clean the data and obtain robust HERs that can be compared across groups or conditions. Our review shows that 55% of the studies (72/132) removed noisy epochs, while 11% (15/132) explicitly mention that bad channels were excluded. The majority of pipelines (73%, 97/133; Fig. 3C) applied independent component analysis (ICA), a commonly used decomposition method designed to separate the recorded M/EEG signal into its underlying brain and non-brain sources.

The primary artifact sources that are addressed by ICA include eye movement (89%, 85 out of 96 pipelines that reported the types of rejected components), cardiac (72%, 69/96), muscle (23%, 22/96), or other nuisance sources (Fig. 3D). While the removal of cardiac artifacts is discussed in Section 3.5, it is important to recognize that other physiological activity such as eye movements (Ohl et al., 2016), muscle activity (Niizeki et al., 1993), or respiration (Yasuma & Hayano, 2004) also co-vary with the heartbeat. The extent to which these factors influence the measured HERs, and whether ICA can mitigate their effects, remains an open question that should be addressed in future studies.

### 3.4. ECG acquisition and preprocessing

ECG recordings are primarily used to extract cardiac markers for time-locking the HER (typically, either R-peak or T-peak). At the same time, they are also often (by 59/132 studies, 45%) employed in control analyses to ensure that observed HER effects are not accompanied and thus cannot be explained by differences in cardiac activity. Placement of the ECG electrodes can play an important role in such analyses, as the presence of differences in ECG between experimental conditions has been shown to depend on the ECG derivation of choice (Gray et al., 2007). Moreover, using just one ECG lead to control for the spread of CFA might not be sufficient to capture the three-dimensional CFA in full (Pérez et al., 2005). With these motivations in mind, we evaluated the consistency of ECG electrode setups used in the literature.

Most studies (67%, 89/132) dedicate two electrodes to record ECG (Fig. 4A), but the placement of the electrodes on the body varies substantially. Although lead II is used most frequently (by 36% of studies, 48/132), some studies also use lead I (11%, 15/132) and lead III (2.3%, 3/132), or locations that do not correspond to any of the standard leads (27%, 35/132; marked as ‘N/C’ in Fig. 4B). Even with standard lead configurations, electrode locations vary (e.g., torso vs. limb placement; Fig. 4C) though the significance of this variability remains unclear and may differ between clinical and non-clinical populations (Sheppard et al., 2011).

**Figure 4.**
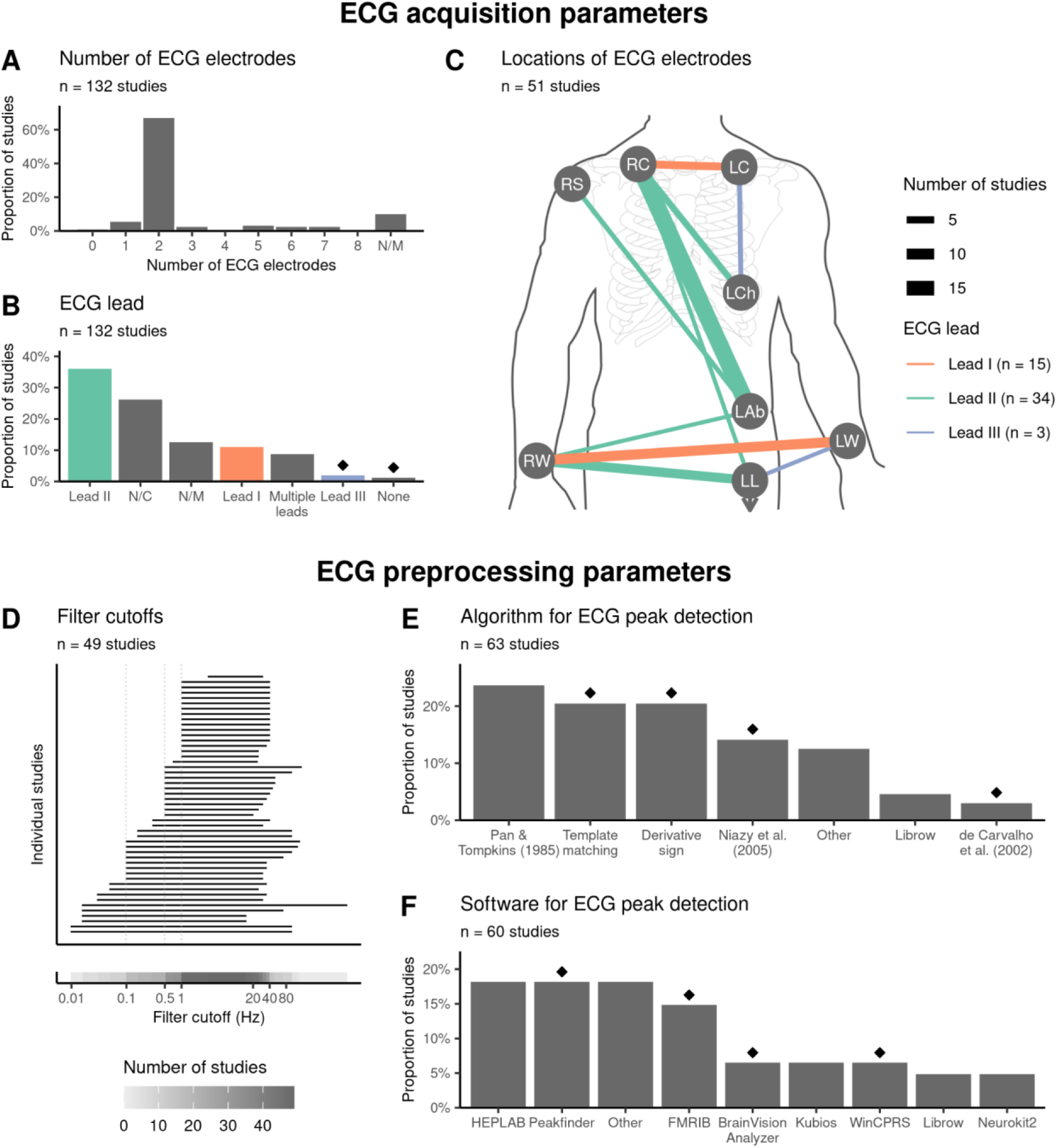
Various ECG leads, electrode locations, and preprocessing settings are used in the literature. Diamond symbols indicate choices driven by one research group (at least 50% of occurrences). (A) Most studies use two electrodes to record ECG. (B) While lead II is the most commonly used derivation, many studies also use electrode placements that do not correspond to any of the standard leads (N/C) or do not specify the lead used (N/M). None denotes studies which reconstructed ECG from EEG. (C) Locations of ECG electrodes that are placed to measure leads I, II, and III. (D) The filtering cutoffs for ECG largely mirror the ones used for M/EEG during preprocessing. (E) Peak detection algorithms based on a template cardiac cycle or ECG derivative are used most often. Category ‘Other’ includes methods that appear only in 1 study. (F) Diverse toolboxes are used for ECG peak detection. Category ‘Other’ includes toolboxes that appear only in 2 studies or less. Abbreviations: RC – right clavicle, LC – left clavicle, RS – right shoulder, LCh – left side of the chest, LAb – lower left part of the abdomen, RW – right wrist, LW – left wrist, LL – left leg/ankle, N/M – not mentioned, N/C – not classified.

In addition, we extracted information about ECG preprocessing settings, including filtering cutoffs and methods for detection of reference events (R-/T-peak). Across the studies, high-and low-pass filter cutoffs largely mirror those used in M/EEG preprocessing, with the most common high-pass cutoffs being 0.1, 0.5, and 1 Hz (Fig. 4D). Especially if researchers plan to use the ECG in statistical control analyses, it may be advisable to match the filter settings across M/EEG and ECG.

Peak detection is typically performed automatically and often followed by visual inspection of the results. As shown in Fig. 4E and 4F, the choice of the method and toolbox for peak detection is highly dependent on the research group. Most common algorithms can be grouped into three major families based on their core principle and target features (Fig. 4E): thresholding of the ECG derivative (Pan & Tompkins, 1985; 15/63, 24%; Niazy et al., 2005; 9/63, 14%), analysis of changes in the derivative’s sign (13/63, 21%), and convolution with a template cardiac cycle (13/63, 21%). As shown in Fig. 4F, commonly used toolboxes include EEGLAB plug-ins HEPLAB (Perakakis, 2019; 11/60, 18%) and FMRIB (Niazy et al., 2005; 9/60, 15%), Peakfinder function (Yoder, 2026; 11/60, 18%), or standalone software, e.g., Kubios (Tarvainen et al., 2014). Notably, peak detection tools might automatically perform data processing steps (e.g., filtering) that are often not explicitly reported, potentially hampering reproducibility.

### 3.5. Dealing with cardiac artifacts

Since HERs are time-locked to the cardiac cycle, it is essential to consider the confounding influence of cardio-vascular artifacts on HER effects. The two major factors that contribute to these artifacts are the electric field generated by the heart (Dirlich et al., 1997) and pulse-related movement of tissue or EEG electrodes (Cataldi et al., 2025; Kern et al., 2013). As mentioned above, researchers predominantly employ one of three overarching strategies to mitigate those effects: avoiding HER space-time points where the influence of artifacts is potentially present, cleaning the data from artifacts, or including the ECG in the statistical analyses. This pattern also held in the reviewed sample of studies, with 89% (117/132) applying at least one of those strategies. However, the applied strategies and their effect on HER might differ substantially.

In terms of artifact removal, ICA is used by 52% of all studies (69/132) to remove cardiac components. This trend is corroborated by research that demonstrates the effectiveness of ICA in attenuating the influence of cardiac artifacts on the measured HERs (Buot et al., 2021). However, there is no standard way of performing ICA, and approaches vary across studies. Most frequently, ICA is applied to continuous data (76%, 74/97 studies that applied ICA). In contrast, around 24% of studies (23/97) choose to perform it after data epoching, likely to specifically target artifacts occurring around the R-peak and T-peak (Dirlich et al., 1997).

While cardiac-related activity often displays relatively consistent spatial patterns across participants (Dirlich et al., 1997), ICA may not always accurately differentiate those from genuine brain signals. Variability of the ICA decomposition across participants can introduce additional variance in the data (Pontifex et al., 2017), potentially influencing HER findings. Given that 86% of the studies that remove cardiac ICs (59/69) also report the approach used for identifying these components, the field is moving towards transparent reporting, facilitating the reproducible removal of cardiac artifacts.

Among the reports, we find that manual identification of CFA components (37/69, 54%) is used more often than automatic (13/69, 19%) or semi-automatic (9/69, 13%) detection approaches (Fig. 5A). The CFA-related components are commonly identified based on the time course (32/69, 46%), topography (31/69, 45%), and power spectrum (13/69, 19%; Fig. 5B). The fully automatic approaches often rely on algorithms such as CORRMAP (Campos Viola et al., 2009), ICLabel (Pion-Tonachini et al., 2019), and SASICA (Chaumon et al., 2015). Semi-automatic cardiac artifact removal is often guided by measures such as phase consistency (7/69, 10%) or correlation (2/69, 2.9%) between ICs and ECG.

**Figure 5.**
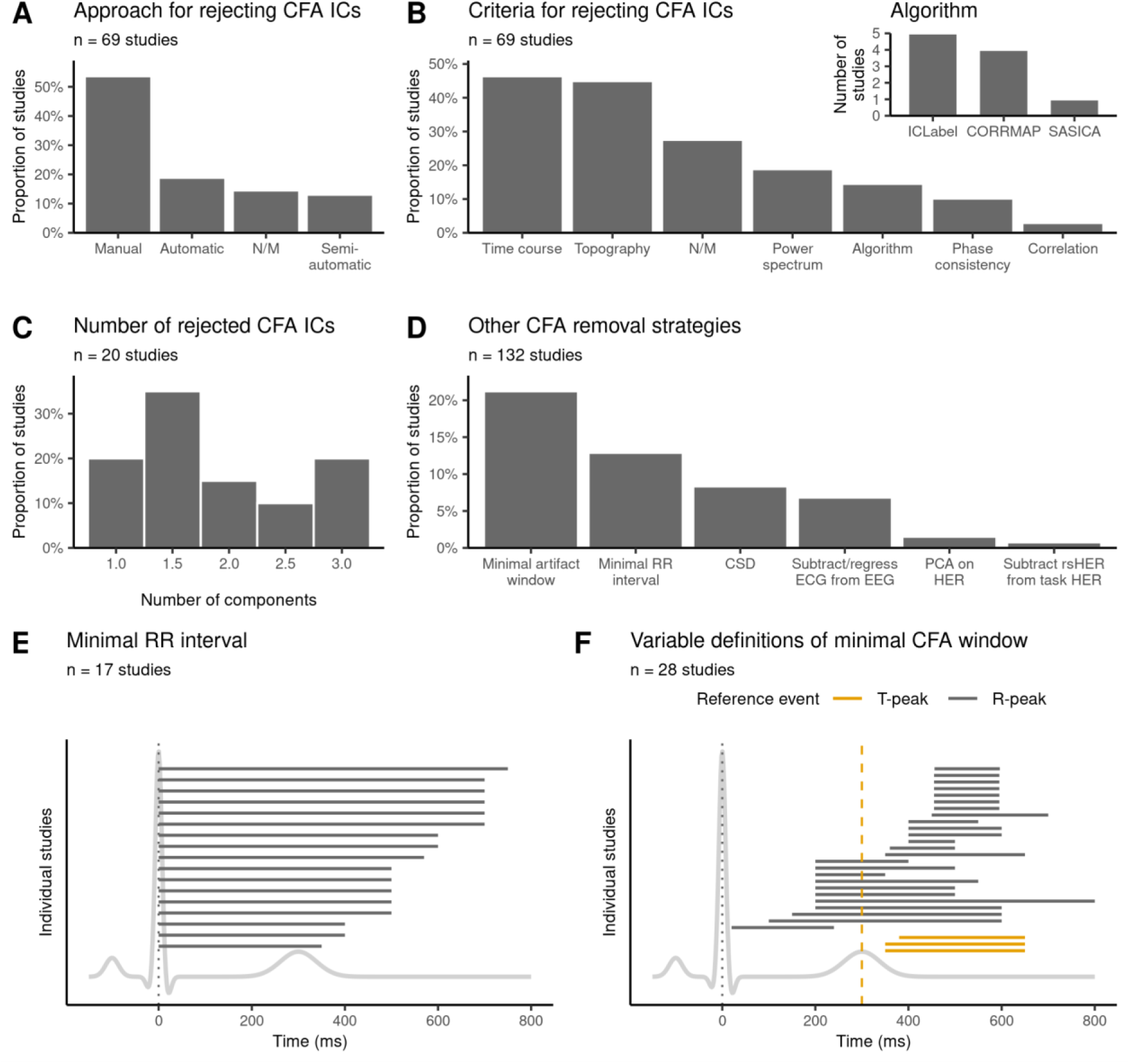
Approaches to address cardiac field artifacts. (A) CFA-related independent components (ICs) are rejected manually, based on automatic approaches or a combination of those (semi-automatic). (B) Decision criteria that are used to define CFA-related ICs. The inset plot displays the number of studies using different algorithms for CFA IC removal. (C) The number of rejected CFA-related ICs per participant ranges from one to three across studies. (D) Additional approaches to address cardiac artifacts. Across the reviewed studies, overlap with artifacts is reduced by (E) excluding trials with short RR intervals, or (F) by using specific time periods of interest for analysis. Simulated ECG traces are shown in gray for reference in panels E and F. T-peak-based time windows (in orange) are shifted by 300 ms relative to the R-peak for illustrative purposes. Abbreviations: ICA – independent component analysis, ICs – independent components, CFA – cardiac field artifact, CSD – current source density, PCA – principal component analysis, rsHER – resting state HER, N/M – not mentioned.

Only 29% of studies (20/69) that removed CFA-related ICs reported the number of removed components, which ranged between 1 and 3 with a median of 1.7 (Fig. 5C). Since many factors, such as the SNR or the M/EEG sensor configuration, influence the number of cardiac ICs obtained by the ICA, it could also be informative to report the combined explained variance of removed cardiac ICs. The explained variance might be a more generalizable measure across studies, allowing for better comparison of CFA removal efficiency. Thus, we suggest considering it for reporting in HER studies (Box 1). Note that differences in cardiac parameters and cleaning may induce spurious HER differences between groups or conditions; it is important to compare the cleaning (number of removed components, explained variance), the number of remaining epochs, and the results without the CFA removal.

Besides ICA, the reviewed studies employ several other techniques to suppress CFA (Fig. 5D). These include current source density analysis (CSD, 11/132, 8.3%), subtracting or regressing out the ECG artifact from the M/EEG data (9/132, 6.8%), principal component analysis (PCA, 2/132, 1.5%), and subtracting the average resting state HERs from task HERs (1/132, 0.8%). However, it remains an open question as to how these different methods compare with one another.

In addition, 21% of studies (28/132) aim to avoid the influence of cardiac artifacts by restricting the time range of interest used in their analyses. Some studies pursue this by excluding early time periods within the cardiac cycle (0-200 ms after the R-peak) from their analyses (Callara et al., 2023; Cambi et al., 2024; Engelen, Buot, et al., 2023; Giusti et al., 2024; Immanuel et al., 2014; Melo et al., 2022). Another common strategy is to choose a time window of approximately 455-595 ms post-R-peak (Gray et al., 2007; Lutz et al., 2019; Müller et al., 2015; Rapp et al., 2023; Schmitz et al., 2020, 2021; Schulz et al., 2013), following Dirlich et al. (1997), who demonstrated the minimal influence of the heart’s electric field during this period. However, given the large variability across studies, there does not appear to be a consensus on a time range free from cardiac artifacts (Fig. 5F).

If HER trials are close to the surrounding heartbeats, it is reasonable to remove such trials to avoid contamination with surrounding HERs or heartbeat-related artifacts. Overall, 13% of studies (17/132) reported the removal of HER trials based on a minimal RR interval, the shortest time allowed between two consecutive R-peaks. We observe that the minimal allowed RR interval varies from 350 to 750 ms across these studies (Fig. 5E). Such large variability could be attributed to diverse participant populations and experimental designs, warranting study-specific definitions of the cutoff. Note that a high minimal RR interval threshold may impose unintended imbalance across the numbers of epochs among conditions or groups if there are underlying differences in heart rate. On top of that, such differences may signify distinct arousal states and distort the interpretation of HER findings. Therefore, it is advisable to report the number of rejected epochs per group and condition (Box 1).

### 3.6. Epoching and baseline correction

Critical decisions in HER research include the choice of the cardiac reference event for epoching (e.g. R-peak vs. T-peak) and the spatiotemporal window used for statistical analysis. These choices can directly influence the detection of HER effects and may define susceptibility to the influence of artifacts. In this review, we find that the vast majority of reviewed studies use the R-peak as reference (123/132, 93.2%). In comparison, only 4.5% (6/132) used the T-peak, and 2.3% (3/132) performed analyses using both peaks within a single study (Fig. 6A).

**Figure 6.**
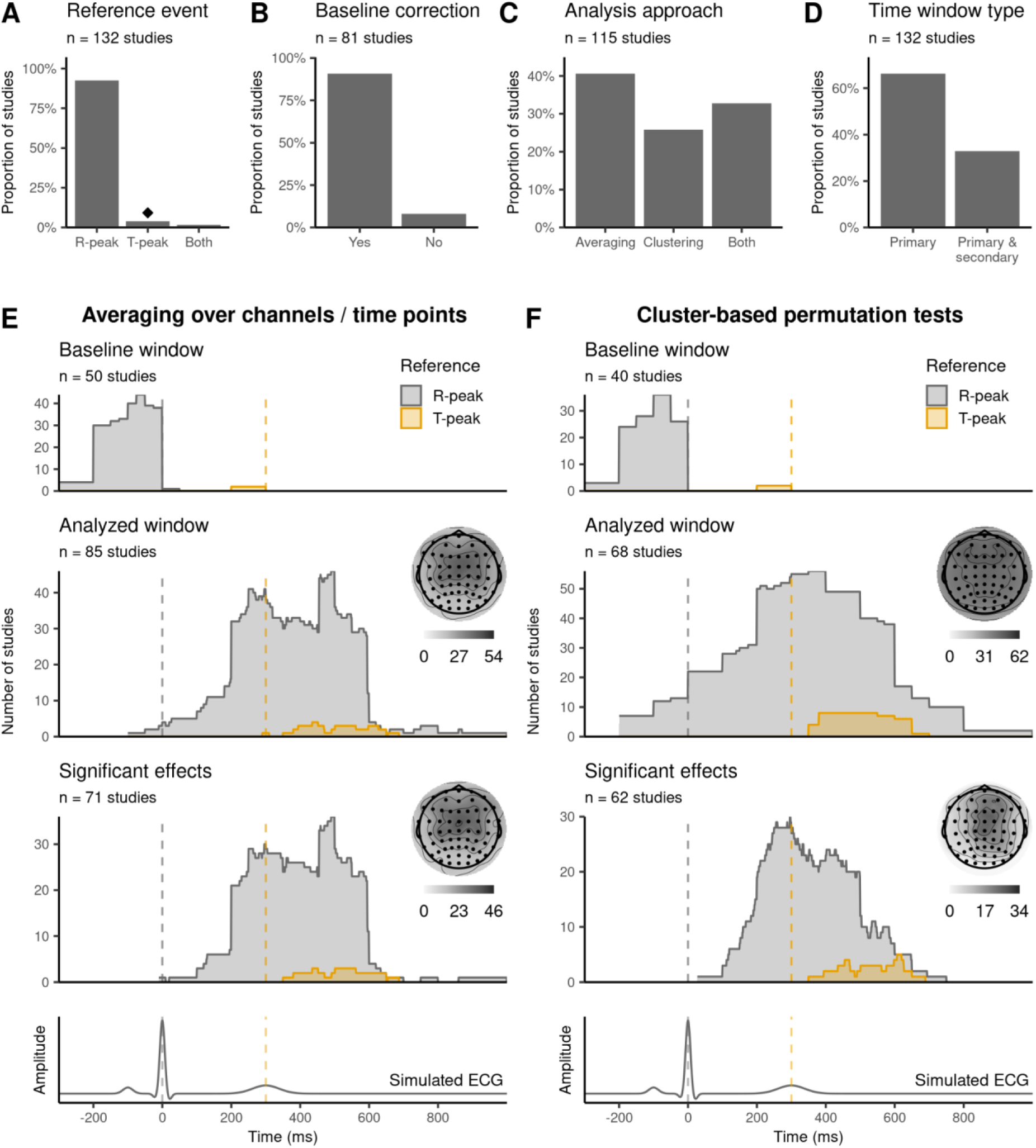
Predefined spatiotemporal regions and detected HER effects. Proportion of studies using: (A) R-vs. T-peak as reference event, (B) baseline correction (applied vs. not applied, unreported cases omitted), (C) averaging approaches vs. cluster-based permutation tests (or both, if performed alongside each other), (D) only primary or both primary and secondary analyses. For (E) averaging approaches or (F) cluster-based permutation tests, baseline windows (top row), time ranges and regions of interest (second row), as well as spatiotemporal distributions of significant effects (third row) are summarized. Panels E and F use cumulative sum plots to visualize the different HER window types, pooled across EEG and MEG studies. Results are split by heartbeat reference R-peak (gray) or T-peak (orange) and aligned to the respective time range of the simulated ECG trace in the bottom row (gray / orange dashed line). T-peak based time windows are shifted by 300 ms relative to the R-peak for illustrative purposes. Topographies include only EEG channels, and colorbars below them indicate the number of studies. Diamond symbol indicates choices driven by one research group (at least 50% of occurrences).

After epoching, baseline correction is typically performed to remove vertical amplitude offsets and slow drifts by subtracting the average pre-stimulus amplitude from the whole epoch. Although it is a standard procedure in conventional ERP analyses (Luck, 2014), it is controversial in HER research, because the heartbeat has a short inter-beat interval (∼1s), potentially allowing activity from one cycle to affect the pre-stimulus time window of the next cardiac cycle (Azzalini et al., 2019).

Overall, only 61% of studies (81/132) clearly state whether baseline correction was performed. Out of these, 91% (74/81) include baseline correction and 8.6% (7/81) state that they do not use it (Fig. 6B). The remaining 39% (51/132) do not mention it in their manuscript, leaving it unclear if they performed baseline correction or just omitted details on this step. Of note, none of the reviewed studies have directly compared baseline-corrected to uncorrected values. Such a comparison could be of value to ensure the observed effects are not due to baseline differences (Box 1). In particular, additional tests may be performed on the baseline window itself or, when performing cluster-based permutation tests, the analysis should be run on the whole epoch, including the baseline.

While 58% of the studies using the R-peak as a reference event (73/126) clearly state the use of baseline correction, only two out of the nine studies using the T-peak (22%) did so. In both studies, the baseline interval was set to -100 and 0 ms relative to the T-peak. A possible explanation for the lower usage of baseline correction, here, could be that the time period preceding the T-peak is close to the cardiac contraction, and hence more prone to contamination by cardiac artifacts.

In R-peak locked analyses, baseline windows show considerable variability, ranging from -300 (min) to 50 (max) ms relative to the R-peak. Notably, 23% of these studies (17/73) define the end of the baseline period at least 25 ms before the R-peak, potentially to avoid including cardiac field artifacts (Coll et al., 2021). One study used a regression-based baseline correction approach to avoid introducing baseline differences in the HER time window of interest (Zaccaro et al., 2024). In future studies, it would be valuable to investigate the effectiveness of baseline-correction methods and contrast them with the caveats associated with not performing any baseline correction.

### 3.7. Predefined spatiotemporal HER regions and resulting findings

When it comes to quantifying the HER signal, amplitude is by far the most prevalent outcome measure (128/132, 97% of studies). Alternative measures are rare, with only three studies using HER latency (2.3%) and two studies examining global field power (1.5%). Note that studies investigating heartbeat-locked neural activity using time-frequency analysis (e.g., Kim & Jeong, 2019) were not considered in this review.

When it comes to choosing regions and time windows of interest, we observe that, overall, analyses locked to T-peak cover a smaller time range (from -10 to 400 ms after T-peak). In comparison, R-peak locked analyses start as early as -250 ms before the R-peak and range until 1000 ms thereafter. Since studies often use averaging over a space-time window of interest or cluster-based permutation tests to identify HER effects (Fig. 6C, studies using only averaging: 47/115, 41%; only clustering: 30/115, 26%; both approaches: 38/115, 33%), we analyzed the data separately for each of these two approaches.

The regions (analyzed for EEG only) and time windows of interest (pooled across EEG and MEG) differ substantially between cluster- and averaging-based analysis approaches. As expected, we see that all EEG channels are typically included in the cluster-based analysis (Fig. 6F). However, averaging is performed more often only over frontocentral EEG channels (Fig. 6E), often motivated by the results of previous studies (Billeci et al., 2021; Elkommos et al., 2023; Schulz et al., 2020). Similarly, clustering approaches use wider time windows more often than averaging approaches do. Nonetheless, there is a clear trend across both categories, averaging as well as clustering, to limit analysis to a time range between 200-600 ms relative to the R-peak (Fig. 6E-F). In MEG studies, it is also common to restrict sensor-space analysis to magnetometers only (Table S3).

Additionally, we also extracted information about channels and time points that show statistically significant test results and pooled them across all reviewed studies. Overall, significant HER effects were found between -10 and 1000 ms for R-peak locked analyses and between 50 and 389 ms for T-peak locked analyses. Interestingly, there is a difference between averaging- and cluster-based analysis in terms of where in time effects are most often detected, as illustrated in the cumulative sum plots in Fig. 6E-F. Specifically for the R-peak locked analyses, comparison of cumulative sum scores indicated that cluster-based approaches identified the most common latencies significantly earlier than averaging (mode = 250 ms vs. 490 ms; permutation test on the time difference between peaks, *p* = 0.021; 1000 permutations). Although the reason for this discrepancy is unclear, one possibility is that cluster-based permutation tests capture effects that are outside of narrow averaging windows by scanning a wider time range. Another possibility is that shorter effects are diluted when averaged across wider time ranges. In contrast, T-peak locked analyses showed no significant difference between clustering (mode = 307 ms) and averaging (mode = 234 ms; permutation test on time difference between peaks, *p* = 0.744; 1000 permutations), likely related to the overall smaller time range of interest in T-peak locked analyses. However, it is important to note that permutation clustering does not allow for any precise inference about the spatiotemporal localization of effects (Sassenhagen & Draschkow, 2019), which means that the comparison to averaging-based analysis should be interpreted with caution. Also note that between-participant variability in R-T intervals may affect the presence or distribution of HER effects in R- or T-peak locked analyses, so it may be worthwhile to compare such analyses. However, few reviewed studies did so (2.3%, 3/132).

When performing analysis on high-dimensional data, one way to alleviate the multiple comparisons problem is to define spatiotemporal regions of interest in advance (Luck, 2014). Our review indicates that most studies follow this procedure and have predefined analysis windows (88/132, 67%, referred to as primary in Fig. 6D). However, we also observe that in 33% of the studies (44/132), researchers perform a two-step procedure, where the selected space-time window is based on the analyzed data within the same study (referred to as secondary in Fig. 6D). While this approach can be useful to identify the electrode sites and time windows where an effect appears in the studied population (Luck & Gaspelin, 2017), it is not always trivial to prevent the analyses from becoming circular (Kriegeskorte et al., 2010; Sassenhagen & Draschkow, 2019).

### 3.8. Statistical analysis and power considerations

Neuroscientific studies are often underpowered, resulting in low research reliability and a potential waste of resources (Button et al., 2013). To assess power in the growing field of HER research, we estimate the minimal effect sizes that the reviewed pipelines can detect with a power of 80% at a 5% significance level, based on their sample size, number of groups and conditions, as well as the type of chosen statistical test.

Across studies, the number of included participants lies between 1 and 552, with a median of 32 (Fig. 7A), the number of studied groups lies between 1 and 6 (Fig. 7B), and the number of conditions lies between 1 and 20 (Fig. 7C). Many pipelines compare groups or conditions (301/410, 73%), or perform a correlational/regression analysis to study a linear association (109/410, 27%). As mentioned earlier, and where possible (Section 2.3), we approximate the number of heartbeat-related epochs used to obtain HERs for the many studies (73/132, 55%) that did not report this information. The estimated and reported numbers of heartbeat-locked epochs range from 22 to 5517 with a median of 300 averaged epochs per participant and condition (Fig. 7D). As shown in Fig. 7E, the t-test family (one-sample, two-sample) is the predominant choice in statistical analysis (74/222 pipelines, 33%), closely followed by variations of correlation (62/222, 28%) and ANOVA (52/222, 23%).

**Figure 7.**
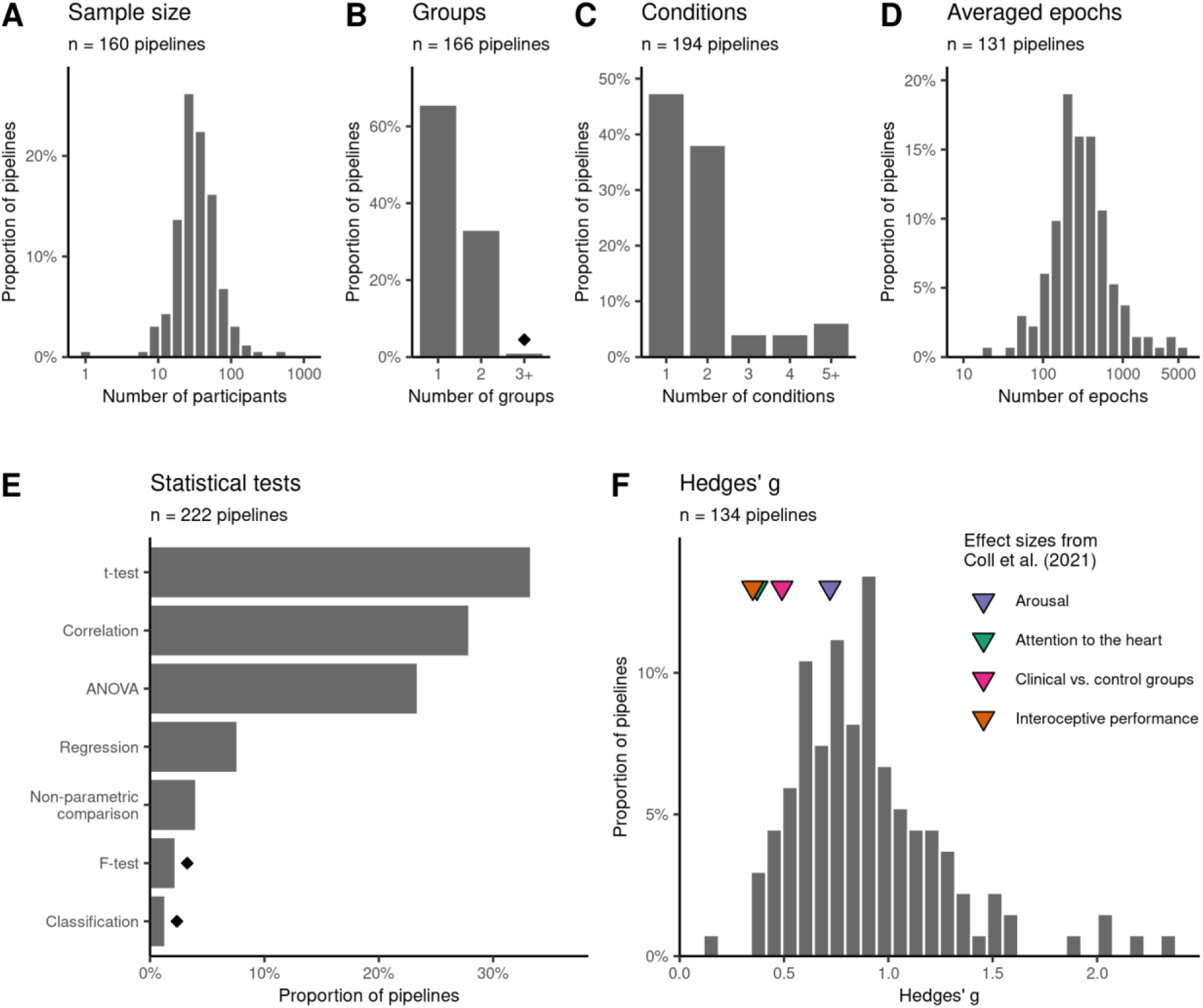
Statistical analysis and minimal detectable effect sizes. The settings used in experimental design and statistical analysis across studies are as follows: (A) sample size (x-axis is log-scaled); (B) number of groups; (C) number of conditions; and (D) estimated number of epochs averaged to obtain HERs per participant and condition (x-axis is log-scaled). (E) The statistical tests that were used in the reviewed studies. (F) Minimal detectable effect sizes with 80% power at 5% significance level, and four HER effect sizes from the meta-analysis by Coll et al. (2021), visualized by colored triangles. Estimated minimal effect sizes are based on the type of statistical test, sample size, number of participants, groups, and conditions, and are only shown for pipelines that use averaged HER values for t-tests, correlations, and ANOVAs. In Panels B and E, diamond symbols indicate choices driven by one research group (at least 50% of occurrences).

The resulting minimal detectable effect sizes range from 0.18 to 2.37, with a median of 0.86 Hedges’ g. Fig. 7F shows their distribution across pipelines that used averaged HER values in one- or two-sample t-tests, correlations, or ANOVAs (estimates including clustering and regression pipelines, which require more assumptions, are shown in Fig. S1). In general, the smaller the minimal detectable effect size, the broader the range of effect sizes that the pipeline can robustly detect. To contextualize these minimal detectable effect sizes, we compared them to meta-level effect sizes from a HER review by Coll et al. (2021). This review investigated the frequently proposed role of HERs as markers of interoceptive functions (Pollatos & Schandry, 2004). The meta-analyses were grouped into categories by topic and assessed whether HER amplitudes are influenced by manipulations of attention (Attention category) or arousal (Arousal category), whether HERs are related to measures of cardiac interoception (Performance category), or whether they differ in clinical populations compared to controls (Clinical category). When comparing the estimated detectable effect sizes to these four meta-level effect sizes, we observe that only 0.7% (1/134) of pipelines had enough power to detect Hedges’ g of 0.35 (Performance meta-estimate), 1.5% (2/134) of pipelines could reliably detect Hedges’ g of 0.37 (Attention meta-estimate), 8.2% (11/134) of pipelines could reliably detect Hedges’ g of 0.49 (Clinical meta-estimate), and 33% (44/134) of pipelines could reliably detect Hedges’ g of 0.72 (Arousal meta-estimate). These observations suggest that 67% (90/134) to 99% (133/134) of pipelines may have yielded underpowered and potentially unreliable findings. Such evidence draws attention to potential reproducibility and reliability problems of current findings within the HER field. Studies with larger sample sizes could test the robustness of previous findings, and future studies may consider adjusting their sampling strategy to detect smaller effect sizes more reliably.

It is essential to note three key points. First, the review by Coll et al. (2021) focuses on a subset of 45 HER studies that report interoception-related HER effects. We did not assess the actual effect sizes among the studies we reviewed, as they varied in topics and experimental designs. Second, the Hedges’ g estimates presented here are based on approximations and therefore should be interpreted with caution. Third, as mentioned previously, HERs are susceptible to noise contamination due to their relatively low amplitudes of 0.5-2 μV (Pollatos & Schandry, 2004; Schandry & Weitkunat, 1990). It is unclear how many trials, or heartbeat-locked epochs, are required to obtain a stable evoked response per participant. Previous ERP studies note that there is no straightforward answer, but it is clear that smaller effects (as typically observed in HER amplitudes) generally require larger samples and more epochs to be reliably detected (Boudewyn et al., 2018; Gibney et al., 2020). However, Hedges’ g estimates discussed in the previous paragraphs do not account for the levels of within-participant noise across HER epochs, i.e., for trial averaging. To examine its effect on minimal detectable effect sizes, we varied the hypothetical within- and between-participant variabilities in HER values and used the numbers of averaged HER epochs from the reviewed studies to obtain adjusted minimal detectable effect sizes (Supplementary Section B and Fig. S2). By construction, they are increasing after adjusting for within-participant variations in HER across epochs. Therefore, in Fig. 7F, we demonstrate optimistic estimates, unadjusted for possible measurement noise. Future methodological studies should examine the levels of HER variability across trials and participants and recommend recording settings. In practice, pilot data can be used to assess HER variability and adjust the number of epochs (see Supplementary Section C on some suggestions). At the very least, we recommend reporting the number of averaged HER epochs per participant and condition (Box 1).

A lack of transparency related to sample size calculation is characteristic of evoked response studies in neuroscience (Larson & Carbine, 2017). To confront this, we recommend preregistering the planned HER analysis alongside power calculations (Box 1). Preregistration is a declaration of planned analysis in a public registry, and it reduces the risk of bias (adjusting methods after knowing outcomes), as well as increases transparency by allowing other researchers to judge the reliability of outcomes (Hardwicke & Wagenmakers, 2023). Amongst the reviewed studies, only 3.8% (5/132) mentioned having published a preregistration. Further, there were no registered reports, advanced forms of preregistration in which the methods are reviewed before data collection and a publication is guaranteed regardless of the outcome (Chambers & Tzavella, 2022). It is possible that the high variability in methods (Sections 3.2-3.6) and findings (Section 3.7) prevents HER researchers from preregistering their hypotheses. This problem can be best remedied if, firstly, most research groups working in the HER field preregister their studies by default, and secondly, more research is devoted to the advancement of robust HER analysis methods.

Another potential benefit of preregistrations is overcoming publication bias, the tendency for publications with significant findings to be published more frequently than those with null findings (Thornton & Lee, 2000). There is evidence for publication bias in the HER field: we observe that only 6.8% of studies (9/132) were published with no significant HER results. We suggest that a combination of preregistrations with formal null testing could reduce publication bias and make null findings more visible and reliable (Box 1). Such results would contribute to the structured accumulation of knowledge in the field, as future studies could use this information to refine their approach. Statistical assessment of “absent” effects is possible using Bayesian statistics (Wagenmakers et al., 2018) and frequentist equivalence testing (Lakens et al., 2018). Few studies in the reviewed sample have used Bayesian statistics to assert the absence of effects, and we did not observe any use of equivalence testing or, importantly, any definitions of the smallest effect sizes of interest. Determining the smallest effect size of interest (Lakens et al., 2018) or the region of practical equivalence (Kruschke, 2021) is not only useful for establishing the absence of effects, but also for making sure that a study does not focus on significant effects too small to be reliably replicated (Box 1).

### 3.9. Control analysis

While small effect sizes could pose serious problems for the reproducibility of HER findings, even the presence of strongly significant, “large” effects does not guarantee that the observed HER effect reflects genuine heartbeat-evoked neural activity (Park & Blanke, 2019; Steinfath et al., 2025). Although many strategies exist to avoid or remove cardiac artifacts (discussed in Section 3.5), it is unclear how effective these are. To ensure that HER effects do not simply reflect cardiac artifacts, 65% of the studies (86/132) include ECG and other heartbeat-related controls in their statistical analysis (Fig. 8A). Specifically, it is common to repeat the analysis on ECG data alone or include it as a regressor of no interest in analyses (combined: 59/132, 45% of studies; Fig. 8B) to demonstrate that effects found at the level of M/EEG sensors do not only reflect cardiac field artifacts. Furthermore, many studies control for the length of the RR interval (45/132, 34%), for instance, by including it as a regressor of no interest in their analyses or by comparing the RR intervals of the studied groups. In addition, 26% of studies (34/132) control for heart rate variability, 6.1% (8/132) for respiration, and 3% (4/132) for blood pressure (Fig. 8A).

**Figure 8.**
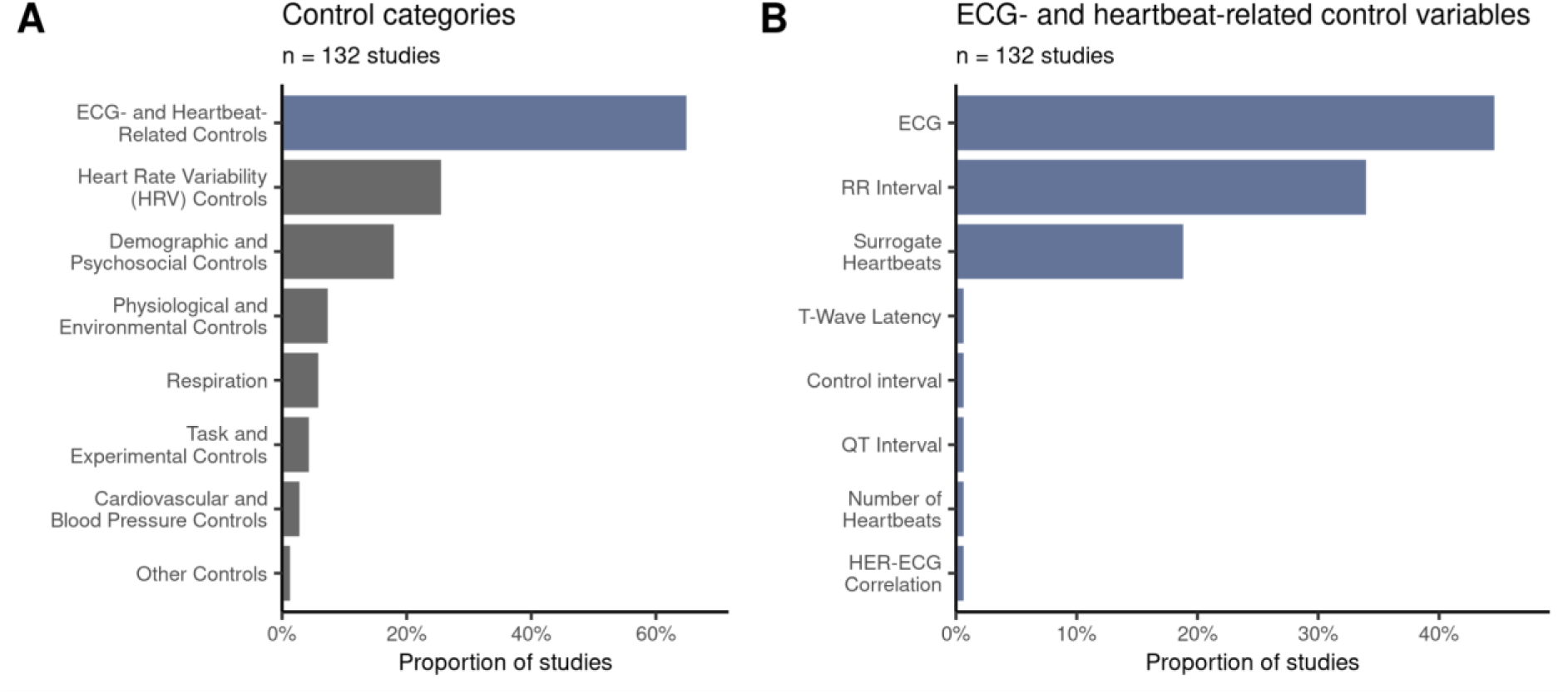
Variables and approaches commonly used for control analysis sorted by the frequency of occurrence in descending order. (A) Control variables grouped by category. (B) Detailed breakdown of ECG- and heartbeat-related control variables. Blue color indicates that items shown in Panel B compose the category ECG and Heartbeat Related Controls from Panel A.

Apart from confounding factors that are directly related to cardiac activity, various heartbeat-independent factors can also affect HER. For instance, if HERs were recorded on-task, the concurrent activity evoked by external stimuli can overlap with HERs, leading to spurious results. Similarly, group-related differences in SNR, for example, due to an imbalanced number of trials, systematic differences in artifacts, comparisons between patients and controls, or differences across age groups, can lead to contaminated HER results. These problems can often be addressed by surrogate heartbeat control analysis or pseudotrial correction (Park et al., 2014; Steinfath et al., 2025). Pseudotrial correction works by subtracting heartbeat-independent confounding activity from the HERs. In contrast, surrogate heartbeat control analyses verify that observed effects are truly locked to the heartbeat by shuffling the onset timings of the cardiac markers within each condition, and repeating the same analyses multiple times (e.g., >100 iterations). The outcome statistic of the original analysis is then compared to the distribution of statistics obtained from these permutations. Hence, it can only be concluded that the original effect is truly related to the heartbeat if it exceeds most of the permuted effects. In our review, we find that 19% of the studies (25/132) perform surrogate heartbeat control analyses or other variations of data shuffling. Note that both methods require a temporal jitter between task and heartbeat onsets. In addition, the way that heartbeats are shuffled within an iteration affects the quality of control analysis. We recommend staying as close to the actual data as possible: same number of epochs, similar IBI and distance to task events, etc., and to clearly describe the motivation for these choices.

Currently, robust control methods for HER research are still being developed (most are listed in the quality control checklist; Box 1). For a full overview of control variables that appeared in the reviewed studies, see Fig. S4.

#### Box 1.

Quality-control checklist for HER analysis and reporting. We suggest key reporting points and recommendations for HER studies based on the extracted information from the reviewed studies and prior experience analyzing HERs.

##### Step 1 – Designing an experiment

A. <u>Preregistration</u>. Supports the reproducibility and reliability of findings, as well as the publication of null results. May incorporate several control analysis pipelines (e.g., with and without baseline correction).
B. <u>Relationship between heartbeats and experimental design</u>. HERs can overlap with task-related activity, and they might co-vary with differences in background activity across groups. Avoid overlaps, if possible, or address them in analysis (see Step 6 – Controls). Exact time locking between heartbeats and task triggers can be difficult to deal with, as correction methods require some jitter.
C. <u>ECG electrode leads, ground, and location</u>. Affect ECG-based control analyses. Report lead configuration used for analysis. Consider recording several leads (e.g. Lead I, II, III, and neck electrodes) to account for the three-dimensional field distribution.
D. <u>Recording of physiological activity.</u> Breathing, eye movements, cardiac and other parameters can confound HER measures.

##### Step 2 – Data preprocessing

A. <u>Filtering</u>. High- and low-pass filter settings can affect the HER shape. High-pass filtering could attenuate genuine HER activity. Hence, for slow HER components, consider a high-pass of 0.5 Hz, which can also diminish respiration-related artifacts.
B. <u>Reference electrode</u>. The choice of online and offline references can affect HER results and artifact removal. Common average reference is widely used and facilitates comparability across studies.
C. <u>R-peak or T-peak.</u> Or other points of reference. Can be compared to each other as they may result in different findings.
D. <u>Minimal IBI</u>. Prevents contamination by overlapping heartbeats. Longer thresholds may induce imbalance in the number of epochs if the heart rate is different between conditions or groups, potentially signifying differences in arousal. Report the number of removed epochs.
E. <u>General cleaning</u>. Report approaches used for noise removal. For ICA, include the description of the algorithm, types of identified components (e.g., ocular, muscle), and rejection criteria. Cleaning algorithms improve reproducibility, but require manual checks.

##### Step 3 – Cardiac Field Artifact Removal (CFA)

A. <u>Rejection of cardiac ICs</u> (if applicable). State whether ICA was applied to epoched or continuous data. ICA on epoched data might improve CFA detectability. Describe rejection criteria, number of removed cardiac-related components, and explained variance.
B. <u>Other CFA removal methods</u>. Detailed description of other approaches used to remove CFA. For example, regressing out ECG from M/EEG data. The HER signal is likely never artifact-free.
C. <u>Limiting analysis time window</u> (optional). Avoid time ranges that are prone to cardiac artifact contamination, e.g., limit analysis to time after T- and before P-wave.

##### Step 4 – HER computation

A. <u>ROI in time and space.</u> Describe start/end times & preselected sensors as well as the motivation for the selection, also for cluster-based permutation tests.
B. <u>Baseline correction</u>. Clearly state if omitted or performed. We recommend avoiding baseline-correction, and testing if the baseline window shows differences between groups or conditions of interest. If performed, include information on the method and time window.
C. <u>Number of epochs</u>. Describe how many epochs were averaged or otherwise analyzed to obtain HER per participant, group, and condition. The number of epochs affects the signal-to-noise ratio, and imbalance may result in spurious findings.
D. <u>Visualization</u>. Show figures with HER results as evoked responses and as topographies.

##### Step 5 – Statistical analysis

A. <u>Statistical power.</u> Ensure that the analysis has sufficient statistical power to detect expected results reliably. E.g., simulations that can be combined with resampling pilot data.
B. <u>Regions of practical equivalence.</u> For both Bayesian and frequentist statistics, define the smallest effect of interest to test if observed effects are negligibly small (i.e., “absent”) and to avoid highlighting practically irreproducible (too small) effects.

##### Step 6 – Controls

A. <u>Surrogate heartbeat analysis</u>. Confirms that observed effects are genuinely locked to the heartbeat but has to be carefully designed. To be valid, they should match real heartbeat properties (e.g. IBI, time to stimuli, number of trials, etc.). Describe how it was performed and how exactly it addressed potential confounding effects.
B. <u>Task overlap controls</u>. For example, pseudotrial correction can help isolate heartbeat-driven effects from task-related activity. Describe how it was performed and how exactly it addressed potential confounding effects.
C. <u>ECG control analysis</u>. Assesses whether differences in ECG exist that might confound the HERs via cardiac artifacts, e.g. by repeating statistical analysis on the ECG channel or regressing out ECG amplitudes. Use multiple leads if possible.
D. <u>Effect of baseline</u>. It is advisable to show the HER results without baseline correction and test if the baseline itself shows significant effects. When performing cluster-based testing, include the baseline time window into analysis.
E. <u>Effect of CFA removal approach</u>. It is advisable to report the HER results without CFA removal to show that they are not explained by different CFA removal efficacy across groups or conditions.
F. <u>Other cardiac parameters.</u> Should be included in statistical analysis. Examples: heart rate, heart rate variability, blood pressure.

##### Step 7 – After publication

A. <u>Data sharing</u>. Publicly shared data enable other researchers to reproduce current findings, develop new research questions, and, importantly, further HER methods. This facilitates more reliable, faster, cost-effective research.
B. <u>Code sharing</u>. Allows for the highest level of transparency in understanding the analysis steps.

Abbreviations: HERs – heartbeat-evoked responses, ECG – electrocardiography, IBI – inter-beat interval, M/EEG – magneto- or electroencephalography, ICA – independent component analysis, ICs – independent components, ROI – region of interest

## 4. Discussion

This systematic review aimed to comprehensively characterize the diverse set of methods used in non-invasive heartbeat-evoked responses research using M/EEG. Our analysis of 132 publications revealed substantial heterogeneity across many processing steps, including data acquisition, data preprocessing, and statistical analyses. While some predominant methodological patterns emerge, such as a higher percentage of EEG compared to MEG studies and the frequent use of the R-peak as a cardiac reference marker, most other steps show considerable variability, as indicated by large entropy scores. However, for the majority of preprocessing steps, studies typically use only one of the many possible options.

Apart from documenting large variability in methodological choices, this review also reveals a large proportion (40-80%) of unreported decisions for processing steps that are critical for HER analysis such as the number of rejected CFA-related independent components (ICs), the number of epochs averaged to obtain HERs per participant and condition, or the baseline window approach used. Taken together with a worrying estimation that 67-99% of studies might be underpowered for detecting meta-level HER effects, it is critical that the community takes more decisive steps towards transparency and reliability of findings.

To alleviate reporting ambiguity and improve the robustness of experimental designs and analyses, we compiled a checklist containing key points that should be considered when conducting and reporting HER studies (Box 1). Since HERs are a subclass of evoked responses, many potential problems are general to the field of M/EEG research and are already summarized elsewhere (e.g. Keil et al., 2014). Furthermore, potential problems relating to the definitions of cardiac phases, a closely related heart-brain topic, are also summarized in a recent review (Caparco et al., 2025). However, some important considerations are unique to HER analysis and require special attention to ensure the quality and reproducibility of results. While previous reviews already highlight some HER-specific best reporting practices (Azzalini et al., 2019; Park & Blanke, 2019), a comprehensive list of reporting guidelines could be useful for experienced and new HER researchers, as well as reviewers of HER studies. Our checklist provides a broad overview of existing approaches for tackling artifacts (mainly, the CFA) and other confounders (e.g., overlap with task-evoked activity).

Despite the overall lack of consensus in methodological approaches, our results highlight the most common choices for each processing step. However, it is important to note that the correctness and suitability of each choice cannot be judged solely based on its occurrence frequency, especially when most studies only use one choice for each processing step. To better understand the impact of the various methodological choices on the observed effects, it is crucial to either perform simulation-based studies with known ground truth derived from single cell recordings (De Falco et al., 2024), intracranial measurements (Kern et al., 2013), or to perform multiverse analyses on real data (Steegen et al., 2016).

Thus, we provide an overview of the currently used methods that could serve as a starting point for novel experimental designs or systematic method comparisons. Based on this overview and previous literature, we have also compiled a tentative HER pipeline intended to provide initial guidance and some unification for HER analysis, rather than to describe strict rules (Figure 9). In addition, we highlight several processing steps which are likely to affect HERs, but their influence has not been systematically studied yet (Box 2). Below, we reiterate the motivation for investigating these steps in more detail.

**Figure 9.**
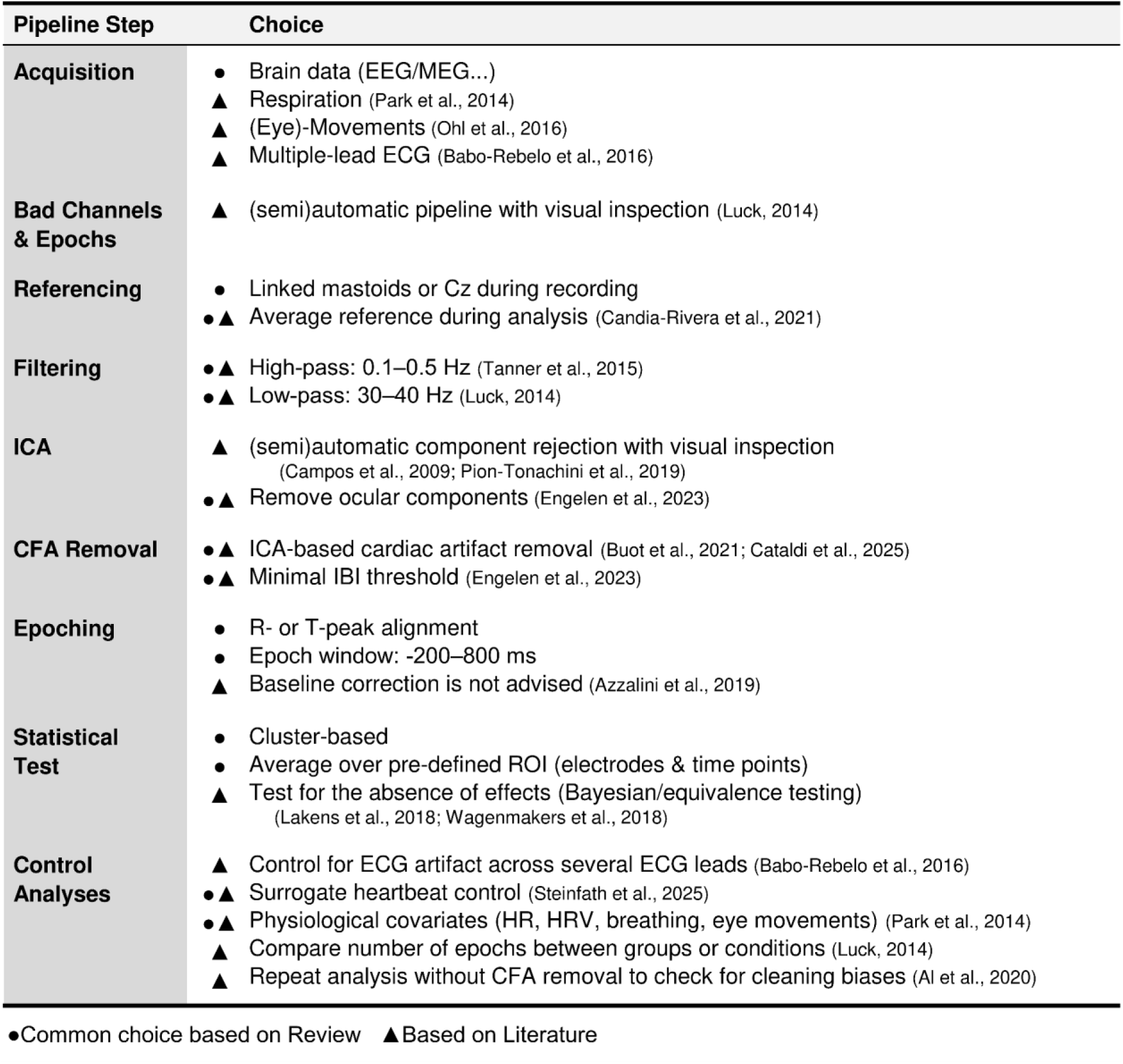
Tentative HER pipeline. Future methodological work is required to determine the best combinations of preprocessing steps and methods for HER research, and the quality of processing choices cannot be judged solely by the frequency of their occurrence in our review. However, to promote comparability and provide practical guidance, we present a tentative pipeline based on suggestions in the literature and common choices among the reviewed studies. Based on the current knowledge, this tentative pipeline can serve as a solid starting point for HER analysis rather than a decisive guideline. See Box 1 for detailed considerations regarding each processing step.

### Box 2.

Outstanding methodological questions in HER research.

To further HER research, the development of methodology is required. Effects of the following processing steps, as well as their interactions, on HER results require further systematic evaluation:

1. High- and low-pass filter parameters
2. ECG recording and subsequent ECG-based CFA controls
3. CFA removal with ICA or other spatial filtering techniques with varying settings
4. Influence of individual differences in cardiac parameters on HER
5. Ocular, muscular, and respiratory artifact influence and cleaning
6. Baseline correction
7. Variability of HER epochs and required number of trials

The high likelihood of being confounded by cardiac artifacts is a major factor that distinguishes HERs from other ERPs. Importantly, the complex three-dimensional nature of cardiac electric fields, with dynamic changes in scalp potential patterns across the heart cycle, poses a critical challenge (Dirlich et al., 1997; Pérez et al., 2005). Decomposition methods, such as ICA, which aim to represent the data in lower-dimensional components, often struggle with time-dependent artifacts. Although the majority of studies use ICA to remove artifacts from the data, there is no consensus on the best approach for identifying CFA-related ICs. While different approaches lead to varying results in artifact suppression, no direct comparison between methods has been conducted to date. It is unlikely, however, that any of the current artifact removal methods result in the perfect suppression of CFA in M/EEG data (Park & Blanke, 2019). Therefore, the HER time courses and topographies observed in the literature represent a complex mixture of residual CFA and neural activity.

A prominent example of this mixture is the deflection in HER epochs reminiscent of the R-peak shape in ECG. It is often observed around 0 ms when the epochs are time-locked to the R-peak. This deflection likely reflects residual CFA in the EEG because it occurs too early to reflect a genuine neural reaction to the R-peak. Interestingly, in some studies, it does not occur exactly at 0 ms. This puzzling observation remains to be resolved in the future, but there may be several hypothetical explanations. Firstly, systematic errors in the identification of R-peaks may result in this shift. It is the responsibility of researchers to ensure that R-peaks are identified precisely. This can be checked by epoching the ECG signal to detected R-peaks, to ensure that the R-peak aligns with 0 ms. Secondly, EEG preprocessing (e.g., downsampling and filtering, CFA cleaning) may lead to these shifts. This can be checked by epoching the raw EEG to detected R-peaks. Thirdly, the shifts may stem from CFA spread and EEG electrode orientation characteristics (Myllylä et al., 2013). Modeling and recording ECG closer to the brain (e.g., neck) might inform this hypothesis.

In addition, groups with differences in cardiac function (e.g., patients vs. controls) might show different contributions of CFA to the M/EEG signal. While this makes artifact removal especially important, it can also complicate the CFA detection and removal. Importantly, cardiac artifact removal itself can induce condition differences. For example, if patients and controls are compared, the ground-truth level of cardiac artifacts might be the same, but due to differences in overall SNR between the groups, CFA removal is more efficient in controls than in patients. Hence, more residual CFA remains in patients, potentially leading to spurious HER differences. One useful control is to verify that results remain overall the same without cardiac artifact correction. However, if results disappear without correction, this does not necessarily indicate that the original findings were driven by inadequate artifact removal, it may also reflect that CFA correction was necessary to reveal the underlying neural signal. A potential proxy for assessing CFA influence on the data is the magnitude of the residual R-peak artifact in the M/EEG recording. Since this artifact results from volume conduction, it likely contains no contribution from neuronal activity. A significant difference in R-peak artifact amplitude between groups or conditions would therefore be cause for concern, as it indicates differential CFA contributions that could confound the HER.

Thus, CFA identification and removal pose a significant challenge. On the one hand, simulations are needed to evaluate the influence and removal efficacy of cardiac artifacts on the measured signal. On the other hand, high-density ECG recordings with sophisticated forward models offer a promising tool for a better separation of CFA and brain activity in real data. The complex nature of the CFA is also affecting ECG-based control analyses, which aim to confirm that HER effects reflect differences in brain activity, rather than volume-conducted CFA. Although low-density ECG derivations are generally sufficient for identifying R- or T-peaks as reference events, they only capture a portion of the complex cardiac electrical field. Since experimental conditions can induce changes in ECG that might be detectable at some recording sites but not at others (Gray et al. 2007), several ECG electrodes covering all three dimensions of the ECG are necessary to show that HER effects are not volume-conducted CFA.

Even though the removal of cardiac artifacts plays an important role in HER research, artifact correction also carries the risk of removing genuine parts from the HER. Hence, many researchers analyze predefined time ranges for their analysis, which show low influence of the CFA (Dirlich et al., 1997). While early HER studies aimed to identify spatiotemporal points where the influence of cardiac-related artifacts is minimal (Dirlich et al., 1997), our review finds little progress in recent years toward identifying and validating these optimal time ranges. Therefore, it remains essential to ensure that changes in HER are not accompanied by changes in cardiac parameters.

Our review supports the idea of considerable temporal variability with predominantly frontocentral HER effects across M/EEG studies. We observe a wide range of chosen time windows, with a preference for limiting analyses to a time range of approximately 200 to 600 ms after the R-peak. This time range is in agreement with intracranial recordings, revealing heartbeat-related brain activity to emerge primarily between 200 and 400 ms after the R-peak (De Falco et al., 2024; Kern et al., 2013). However, even in these invasive studies, there is considerable variability regarding the onset, duration, and anatomical location of cardiac-related neural responses. This spatiotemporal variability in observed effects suggests that there is no unique empirically observed manifestation of HER (Gautier et al., 2025), but that it reflects a mixture of responses to different parts of the cardiac cycle, or even a sustained, ongoing tracking signal related to blood pressure and heart rate variations. Overall, those pieces of evidence point to a wide range of cardiac-locked modulations of neuronal activity depending on the specific task, cognitive state, experimental condition, or pathology.

In line with this, another potential source of variability in observed locations of HER effects that obscures comparability is the time-locking reference for HER epochs. While most studies choose R-peak as a reference (123/132, 93.2%), some use T-peak (6/132, 4.5%), and only a few compare both within a single study (3/132, 2.3%). Due to interindividual variability in R-T intervals, the two approaches may result in different spatiotemporal locations of effects and are not directly comparable. Because only a few studies choose both R- and T-peak locking, it is difficult to judge how often the results would be similar or different and why. Ideally, more studies investigate this, and in the future, more mechanistic explanations are proposed. It would also be informative to see larger HER epochs in more studies: how the effects look when we consider a period spanning several heartbeats, e.g., from -2 to 2 seconds around the R-peak, and how the results depend on the point of reference within the cardiac cycle.

The spatiotemporal variability of the HER affects the suitable analysis options. Due to the inconsistency of results across studies, there is currently no clear spatiotemporal region that shows robust HER effects. Hence, many researchers use hypothesis-free cluster-based permutation testing. While this approach is convenient to circumvent the variable nature of the HER, it is important to keep in mind that cluster-based permutation tests do not allow for exact spatial or temporal interpretation of effects (Maris & Oostenveld, 2007; Sassenhagen & Draschkow, 2019). In addition, we observe that many studies use a two-step procedure, where the region of interest and time range are first defined based on primary cluster-based permutation analysis. The resulting cluster is then used in further secondary analyses. However, if the analysis used for data selection is not independent of the secondary analysis, circularity may become a concern (Kriegeskorte et al., 2009). While this does not mean that the results are incorrect, it underscores the need for independent replication and careful evaluation of the statistical validity of such two-step analyses. Ideally, cluster-based permutation analyses with a more exploratory nature are followed by clear confirmatory studies with predefined regions of interest on separate datasets, allowing for more straightforward power estimation and tests for the absence of effects as well as facilitating replicability. However, in studies that aim to establish the presence of HER differences, e.g., between conditions or groups of patients, without the focus on exact spatiotemporal locations, conventional cluster-based analysis remains a viable option (Rousselet, 2025). Moreover, the cluster-based tests are not the only way to correct for multiple comparisons when performing mass univariate tests on spatiotemporal points in neurophysiological data: other ways include traditional conservative approaches such as Bonferroni (Armstrong, 2014) or Benjamini-Hochberg (Benjamini & Hochberg, 1995) corrections, as well as novel applications of Bayesian generalized additive multilevel models (Nalborczyk & Bürkner, 2025) or cluster-depth tests (Frossard & Renaud, 2022).

Careful experimental design and control analyses are essential to isolate genuine HER activity from other sources of variability. When HERs are studied in task settings, as is the case in most of the reviewed studies, task-related evoked responses may overlap with internal responses like the HER. This can become problematic if, for example, experimental conditions systematically differ in the temporal alignment between heartbeats and task triggers (e.g., presented stimuli), as the resulting HERs might capture different parts of the stimulus-evoked responses, leading to spurious HER differences. This opens additional challenges. On the one hand, researchers have to make sure that potential HER effects are not a result of temporally overlapping task-related differences in neural activity. Currently, we find that there are two approaches used in the literature to deal with this situation: surrogate heartbeat analyses (Park et al., 2014) and pseudotrial correction (Steinfath et al., 2025). On the other hand, if effects in task-evoked activity are observed, researchers have to make sure that these are not a result of overlapping cardiac-related activity, given that heartbeats may align to task triggers (e.g., Kunzendorf et al., 2019). Overall, disentangling overlapping effects might also be approached by regression-based ERP modelling (Ehinger & Dimigen, 2019). However, the studies included in our review indicate that, so far, these methods have not gained wider use in the HER research community. Further work is therefore required to show in which scenarios the different methods work best and where the limitations are. Such limitations include tasks where a tight time-locking between HERs and external stimuli is present (e.g., tones presented synchronously to the heartbeat), because current correction methods depend on jitter to delineate HER activity from task-evoked responses.

Furthermore, the mere presence of surrogate analysis or pseudotrial correction does not guarantee the proper control for potential confounders. Depending on the experimental paradigm and the confounder in question, specific time windows or numbers of simulated epochs need to be carefully chosen. If a group difference in HERs arises from unequal SNR due to imbalanced numbers of trials, it is important that the number of surrogate heartbeats is preserved in the control analysis. Generating surrogate heartbeats with an arbitrary number of epochs would alter the variance and hence the statistical power of the surrogate test, undermining its validity. Likewise, the temporal relationship between HER and confounding factors has to be preserved. For example, if heartbeats occur at different delays relative to task stimuli across conditions, the resulting HER epochs capture different portions of the task-evoked ERP (Steinfath et al., 2025). For the surrogate analysis to properly control for this, the surrogate trials must maintain the temporal distribution of heartbeats relative to the task events within each condition. When the heartbeats are shuffled randomly, without respecting the original timings, the overlap with the task-evoked ERP would be disrupted, thus invalidating the generated null distribution. As the guidelines on the robust design of surrogate control analysis depend on the individual experimental design, we advise that the authors clearly state how exactly they performed it, what problem they were trying to address, and how the surrogate method solved it. This level of transparency enables the readers to make judgments about the quality of surrogate analysis, beyond the fact that it was performed.

Adding to these challenges, the heart does not operate in isolation and is coupled to several other physiological systems (Hall & Hall, 2021). This coupling can complicate the separation of genuine HER effects from other correlated sources of variability, such as eye movements (Ohl et al., 2016) or respiration (Yasuma & Hayano, 2004; Zaccaro et al., 2024). Consequently, careful experimental design and control analyses are required to delineate heartbeat-related effects from other influences. Importantly, our review revealed that, across HER studies, a broad range of different control variables has been included in the analyses. While eye movements or respiration are not among the most frequent choices, some experimental designs may be more prone to these specific confounds, which warrant explicit care. As we did not assess the adequacy of control analyses across the reviewed studies, some results, especially spatiotemporal locations of detected HER effects, are likely to be affected by traces of various artifacts and other sources of activity (task, physiology) and should be interpreted with caution. We encourage the readers to apply stricter exclusion criteria (e.g., presence of ECG or respiration controls and surrogate analysis) in the web application we provide (https://paulsteinfath.shinyapps.io/her-systematic-review/), which offers updated, dynamically generated versions of the figures presented in this review.

The current review has several limitations. Since only information about methods was extracted, we cannot compare and recommend any methods solely based on how often they are used. Furthermore, methods used by particular HER research laboratories might be overrepresented in the overall results. In fact, we observed that 20% of studies in the reviewed sample (27/132) have the same three last authors (max. 12 studies per last author). However, we added a ‘dominance indicator’ represented by a diamond to all histograms to foster transparency. Most of the processing choices we reviewed do not represent research practices of any particular group of authors, except for ECG recording and preprocessing. Other determinants, such as the study sample characteristics (patients or controls, age groups) and research topics, may contribute to the reported variability in methods and findings, but an in-depth analysis of these factors is beyond the scope of this review. However, our interactive resource contains the relevant information for further inquiries. Another limitation is that we did not extract the effect sizes per reviewed study or relate them to the employed methods due to the substantial heterogeneity in research questions. Therefore, an objective comparison of methodological choices for HER analysis on the same dataset is a crucial next step for the advancement of HER methodology. Additionally, we did not extensively extract all non-significant control analyses, which might also bias the representativeness of some of the results. It is critical to remember that any review is subject to publication bias when it comes to the included studies (Thornton & Lee, 2000). Overall, we find that a stunning 93% of reviewed studies reported statistically significant findings. Given that we also observe that a large proportion of studies might be underpowered, the small number of studies with non-significant findings is another evidence of a possible publication bias (Button et al., 2013).

The problem of publication bias, however, can be remedied in the future of the HER field by pursuing more transparent practices such as preregistrations and registered reports, and reporting the significance of null findings through Bayesian approaches (Wagenmakers et al., 2018) or frequentist equivalence testing (Lakens et al., 2018). We suggest that, just as for other research domains, preregistrations and especially registered reports can increase the quality of HER research and are a key solution to many of the issues outlined in this review (Chambers & Tzavella, 2022; Hardwicke & Wagenmakers, 2023; Soderberg et al., 2021). In such a growing field, it might be difficult to navigate the multiplicity of methodological choices across the many processing and analysis steps, and prepare a robust research plan in advance. This is further complicated by variability in reported HER findings. Therefore, based on this review, literature, and our current knowledge, we propose a tentative HER pipeline, which, along with a broader quality-control checklist (Box 1), could serve as a solid starting point for future investigations (Figure 9). Characterizing the temporal, spatial, and spectral components of HER across contexts is the key direction of future research in the field, but this, first and foremost, warrants the advancement of understanding how to analyze HERs in the most robust way and heavily relies on transparent and reliable findings. This review and future, much-needed, methodology-oriented HER studies will shed light on these decisions and help researchers in preparing their HER studies. Ultimately, accumulation of robust evidence through reliable research with transparent methodology is the way to understanding what HER is, and thereby, how the brain and body interact.

Our review provides a set of reasonable candidate methods to consider for the preprocessing and analysis of HERs, and the extracted information is published with the article. To make the results more accessible, we also provide an easy-to-use interactive interface for inspecting the dataset. We hope that this review will serve as a foundation for improving the methodological consistency across HER studies.

## Statement on gender composition within HER research and citation diversity

Recent evidence suggests that publications with female names at prominent positions in the author list are cited less often in the field of neuroscience than should be expected (Dworkin et al., 2020; Hefter et al., 2025). In light of recent discussions by journals and publishers (e.g., “Citation Diversity Statement”, 2023; Fulvio et al., 2021; Sweet, 2021), we analyzed gender approximations across the HER literature included in this review to guide the benchmarking of future citation practices.

As done by Dworkin et al. (2020), gender estimates (F = female; M = male) were extracted for the names of the first and last authors across all references (n = 132) through an automated service (https://gender-api.com) and the cleanBib toolbox (D. Zhou et al., 2022). In case of low assignment probability (<.80), publicly available pronouns were manually searched for. Following the exclusion of those for which this failed (n = 2), the author pairs were classified as 18% (n = 24) FF (first author/last author), 10% (n = 13) MF, 31% (n = 40) FM, and 41% (n = 53) MM. There were no publications by singular authors. Four publications categorized as FM were authored by our lab.

A median split by publication year showed that gender balance improved over time (since October 2020 [Δ before]: 23% FF [+9.2%], 14% MF [+7.7%], 28% FM [-6.2%], 35% MM [-11%]). The observed pattern resembles the gender distribution (18% FF, 14% MF, 29% FM, 39% MM) that resulted from large-scale analyses by Hefter et al. (2025). For more detailed analysis see Section E of the Supplementary Material.

To note, the applied procedure falls short of some aspects: 1) shared first and/or last authorship were not considered, 2) gender estimates and openly accessible pronouns may not have been, in full, reflective of the respective individuals’ gender identities, and 3) gender was taken as dichotomous variable only, and thus, the procedure does not allow for monitoring gender representation in its full variety. Lastly, some clustering of HER publications (using a threshold of ≥5) was observed, with 27 studies (20%) last authored by the same three names (2 F, 1 M), and 15 studies (11%) first authored by another set of three names (1 F, 2 M).

Lastly, we examined our own citation behavior in the main text (n = 111). After excluding four references with undetermined author gender, the remaining were classified as 15% (n = 16) FF, 12% (n = 13) MF, 23% (n = 25) FM, and 50% (n = 53) MM. These proportions include nine single-authored studies (1 FF, 8 MM) and five from our lab (2 MM; 3 FM, with two amongst the reviewed studies). In total, 35 of the 132 studies included in the review (26.5%) were also cited in the main text.

## Studies included in the review

(Al et al., 2020, 2021, 2023; Alkhachroum et al., 2024; Azzalini et al., 2021; Babo-Rebelo et al., 2019; Babo-Rebelo, Richter, et al., 2016; Babo-Rebelo, Wolpert, et al., 2016; Banellis & Cruse, 2020; Baranauskas et al., 2017; Billeci et al., 2021; Birba et al., 2022; Bogdány et al., 2022; Buot et al., 2021; Callara et al., 2023; Cambi et al., 2024; Candia-Rivera, Catrambone, et al., 2021; Canales-Johnson et al., 2015; Candia-Rivera, Annen, et al., 2021; Candia-Rivera et al., 2023; Candia-Rivera & Machado, 2023a, 2023b; Couto et al., 2014, 2015; de la Fuente et al., 2019; Di Bernardi Luft & Bhattacharya, 2015; Dirlich et al., 1997, 1998; Elkommos et al., 2023; Engelen, Buot, et al., 2023; Fittipaldi et al., 2020; Flasbeck et al., 2020, 2021, 2024; Fouragnan et al., 2024; Fukushima et al., 2011; Gao et al., 2023; García-Cordero et al., 2017, 2016; Gentsch et al., 2019; Giusti et al., 2024; Gray et al., 2007; Hermann et al., 2024; Hodossy et al., 2021; Immanuel et al., 2014; Ito et al., 2019; Judah et al., 2018; Kamp et al., 2021, 2023; Katkin et al., 1991; Kato et al., 2020; Khoshnoud et al., 2022, 2024; Kim et al., 2019; Koreki, 2024; Kritzman et al., 2022; Kumral et al., 2022; Lechinger et al., 2015; Legaz et al., 2022; Leopold & Schandry, 2001; Liu et al., 2022, 2023; Liuzzi et al., 2024; Lutz et al., 2019; MacKinnon et al., 2013; Mai et al., 2018; Maister et al., 2017; Majeed et al., 2022; Marshall et al., 2017, 2018, 2019, 2020, 2022; McCraty et al., 2004; Melo et al., 2022; Migeot et al., 2023; Montoya et al., 1993; Müller et al., 2015; Ono et al., 2024; Pang et al., 2019; Park et al., 2014, 2016; Pelentritou et al., 2024; Pérez et al., 2005; Perogamvros et al., 2019; Petzschner et al., 2019; Pollatos et al., 2005, 2016; Pollatos & Schandry, 2004; Poppa et al., 2022; Rapp et al., 2023; Ren et al., 2022; Richter et al., 2021; Richter & Ibáñez, 2021; Salamone et al., 2018, 2020, 2021; Schandry et al., 1986; Schandry & Montoya, 1996; Schandry & Weitkunat, 1990; Schmitz et al., 2020, 2021; Schulz et al., 2013, 2018, 2020; Schulz, Ferreira de Sá, et al., 2015; Schulz, Köster, et al., 2015; Sel et al., 2016; Shao et al., 2011; Simor et al., 2021; Solcà et al., 2020; Stoupi et al., 2024; Tanaka et al., 2023; Terhaar et al., 2012; Todd et al., 2021; Ventura-Bort & Weymar, 2024; Verdonk et al., 2021, 2024; Villena-González et al., 2017, 2023; Wei et al., 2016; Weijs et al., 2023; Yoris et al., 2017, 2018, 2024; Yuan et al., 2007; Zaccaro et al., 2022, 2024; Zhang et al., 2023; H. Zhou et al., 2022; M. Zhou et al., 2024; Zwienenberg et al., 2023)

## CRediT

P.S.; M.A.; N.K.: Conceptualization, Methodology, Software, Validation, Formal analysis, Investigation, Data Curation, Writing – original draft, Writing – review & editing, Visualization, Project administration. T. L.: Data Curation, Writing – review & editing. L.E.: Gender Citation Diversity Statement, Writing – review & editing. V.N.; A.V.: Writing – review & editing, Supervision, Funding acquisition.

## Data and code availability statement

The full data table containing all relevant information extracted from the literature that is used in this systematic review can be accessed at https://osf.io/znrbm. In addition, we provide an interactive web application that allows browsing the dataset: https://paulsteinfath.shinyapps.io/her-systematic-review/. All analysis scripts are available at https://github.com/PaulSteinfath/systematic-hep-review. Citation diversity code along with gender approximation datasets can be accessed at https://github.com/liobaenk/citation-diversity-tracking.

## Acknowledgement

This project was supported by the Max Planck Society. T.L. received funding from the Ministry of Health and Social Security of the Government of Luxembourg. We thank the reviewers for their valuable comments and the authors of reviewed studies for moving forward the HER field.

## Declaration of generative AI in scientific writing

During the preparation of this work the authors used OpenAI chatGPT (GPT-4o), Anthropic Claude 4, and Grammarly in order to proofread the text, and assist during analysis, code generation and documentation. After using these tools, the authors reviewed and edited the content as needed and take full responsibility for the content of the publication.

## Footnotes

1 In the EEG field, these responses are commonly referred to as heartbeat-evoked potentials (HEPs) due to the nature of the electric potentials recorded with EEG. MEG studies, however, refer to them as heartbeat-evoked fields (HEFs) or heartbeat-evoked responses (HERs). Since we include both EEG and MEG studies in this review, we use the more general term ‘response’ (HER) throughout the text.

