## Supplementary Material for "Heartbeat-evoked responses in M/EEG: A systematic review of methods with suggestions for analysis and reporting"

Contents

A. Estimation of minimal detectable Hedges’ g across reviewed pipelines	2

B. Minimal detectable effect sizes depending on the number of averaged epochs and signal variability	3

C. A practical note on the required number of trials	6

Simulations	6

Estimating between-within variability from pilot data	7

D. Other supplementary figures and tables	9

E. Details on evaluation of gender composition within HER research	13

References	13

### A. Estimation of minimal detectable Hedges’ g across reviewed pipelines

Evaluation of the minimal detectable Hedges’ g was performed under the following assumptions:

- We always considered a two-sided alternative hypothesis.
- We assumed equality of groups, so the sample size per group was equal to the sample size divided by the number of groups.
- We have encountered only a couple of uses of Bayesian statistics across all papers. For estimation purposes, we treated all tests as frequentist.
- In ANOVAs, we assumed that the number of cells (unique clusters of observations that we use for power estimation) was equal to the product of the number of groups and the number of conditions. We also assumed that each group performed all conditions.
- For clustering, we assumed it to be at least as powerful as a test on averaged HERs.
- For regression, we assumed a regression with one predictor.

Since the last two assumptions might not hold in practice, Hedges’ g estimates for cluster-based tests and regression are only shown in Fig. S1 but not in Fig. 7F.

For all considered types of statistical tests, we first estimated Cohen’s d:

- For t-tests, we obtained it directly from the power estimation function of pwr package in R.
- For ANOVAs and regressions, we obtained Cohen’s $f$ from the power estimation function and transformed it as follows (Cohen, 1988):

$$d = 2f$$

- For correlations, we obtained the correlation coefficient (denoted as *r* in the equation below) and converted it to Cohen’s d (Cohen, 1988; Ruscio, 2008):

$$d=\frac{2r}{\sqrt{1-r^{2}}}$$

The exact Hedges & Olkin (1985) correction was performed according to the following formula:

$J(\upsilon) = \frac{\Gamma\left( \frac{\upsilon}{2} \right)}{\sqrt{\frac{\upsilon}{2}\Gamma\left( \frac{\upsilon-1}{2} \right)}}$,

where $\upsilon$ is the number of degrees of freedom. Hedges’ g is then obtained by multiplying Cohen’s d and $J(\upsilon)$. For further implementation details, see the code at: <https://github.com/PaulSteinfath/systematic-hep-review/blob/main/functions/analysis/hedges_g.R>

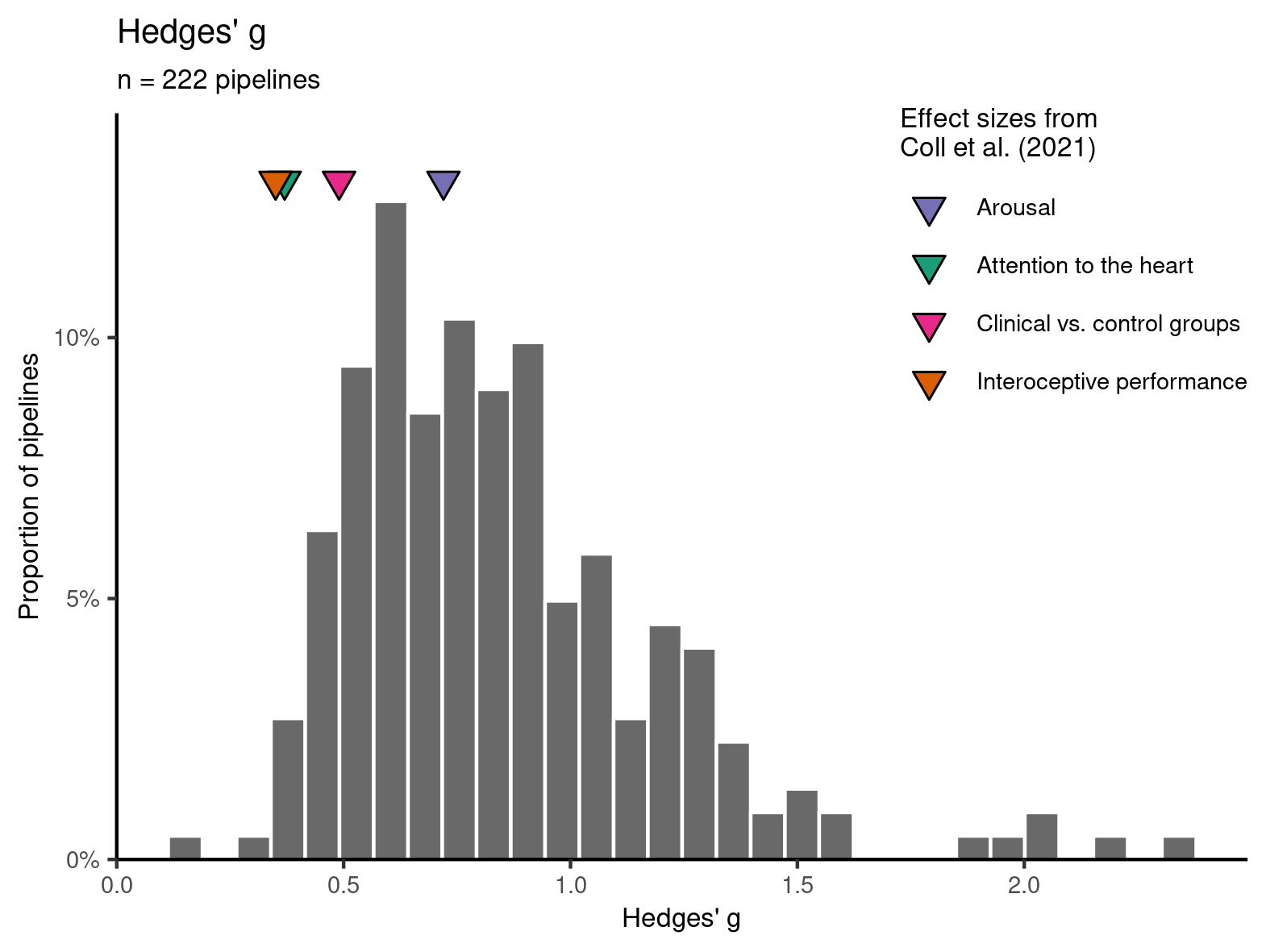

*Figure S1*. Minimal detectable effect sizes across reviewed pipelines. Estimates for cluster-based permutation tests and regression are also included. These additional estimates are not reflected in Fig. 7F, which only includes estimates from one- or two-sample t-tests, ANOVAs, and correlations that were performed on averaged HER values. When comparing the estimated detectable effect sizes to the four effect sizes reported in a previous HER review by Coll et al., 2021 (marked with colored triangles), we observe that only 0.9% of pipelines (2/222) had enough power to detect Hedges’ g of 0.35 (interoceptive performance meta-estimate), 2.3% of pipelines (5/222) could reliably detect Hedges’ g of 0.37 (attention to heart meta-estimate), 9.9% of pipelines (22/222) could reliably detect Hedges’ g of 0.49 (clinical vs. control groups meta-estimate), and 41% of pipelines (91/222) could reliably detect Hedges’ g of 0.72 (arousal meta-estimate).

### B. Minimal detectable effect sizes depending on the number of averaged epochs and signal variability

The estimated effect sizes shown in Fig. 7F and S1 were obtained assuming that we have only one perfect observation per participant or an infinite number of observations, allowing for a perfect estimate. In practice, HERs have relatively low amplitudes (e.g., 0.5–2 μV in the case of EEG), so multiple trials per participant and condition are typically averaged to obtain a more stable response. To understand the influence of the number of trials on the minimal detectable effect size, we varied the levels of between- and within-participant variability in HER values and used the numbers of HER epochs from the reviewed studies to obtain minimal detectable effect sizes adjusted for the number of averaged epochs, according to the formula derived below.

In the following, we consider effect sizes based on standardized mean differences (SMD). This derivation is applicable to a class of frequentist statistical tests that are based on SMD, including (1) one-sample t-tests, (2) paired t-tests, (3) two-sample t-tests (independent groups), (4) simple ANOVA contrasts or linear regressions with standardized coefficients that are algebraically equivalent to SMD (e.g., a standardized regression coefficient on a factor variable with two levels).

Cohen’d is defined as:

$$d=\frac{\mu_{1}-\mu_{2}}{p}$$

where $\mu_{1,2}$ are the means of two groups or conditions (note that $\mu_{2}$ can be 0, which covers the standard one-sample test) and $p$ is the pooled SD (Cohen, 1988). $d$ is the “perfect”, true effect size - the one that does not depend on the number of averaged observations.

We simplify by assuming that (1) for two-sample t-tests, group sizes and between-participant variances are equal; and (2) for paired t-tests, between-participant variances are equal for both conditions and no covariation between conditions exists.

Then, for the case of no averaging over trials and a two-sample or simple one-sample t-test, pooled SD $p$ is equal to the between-participant SD $\sigma_{b}$:

$$p=\sigma_{b}$$

For the paired test, $p$ is equal to the between-participant SD $\sigma_{b\Delta}$ of the change score between two conditions that can also be expressed as $\sigma_{b}\sqrt{2}$.

If we now consider the case when each observation is obtained by averaging $k$ trials per participant (assuming exactly the same number $k$ for each participant and also across groups or conditions, i.e., $k$ trials per condition), we need to take the standard error of the mean into account. Assuming that within-participant SD $\sigma_{w}$ is the same for all participants (independent and identically distributed, also across conditions or groups, if applicable), pooled variance is equal to the sum of between- and within-subject variances. Hence, pooled SD for the case of averaging over $k$ trials is equal to:

$p_{k}=\sqrt{\sigma_{b}^{2}+\frac{\sigma_{w}^{2}}{k}}=\sigma_{b}\sqrt{1+\frac{1}{r^{2}k}}$ where $r=\frac{\sigma_{b}}{\sigma_{w}}$

is the ratio of between- and within-participant SDs (later referred to as between-within variability ratio). Or, in the case of the paired test, similarly expressed as $\sqrt{{2\sigma}_{b}^{2}+2\frac{\sigma_{w}^{2}}{k}}$ .

As a result, the observed effect size with $k$ trials per participant is equal to:

$$d_{obs}=\frac{\mu_{1}-\mu_{2}}{p_{k}}=\frac{\mu_{1}-\mu_{2}}{\sigma_{b}\sqrt{1+\frac{1}{r^{2}k}}}=\frac{d}{\sqrt{1+\frac{1}{r^{2}k}}}$$

Which shows that the effect size adjusted for number of epochs (the observed effect size) is the true effect size multiplied by $\frac{1}{\sqrt{1+\frac{1}{r^{2}k}}}.$ Since the square root in the denominator is always at least 1, the estimated effect size $d_{obs}$ is always smaller than or equal to the true effect size $d$. Equality is reached only in case of infinite $r$ (equivalent to zero within-participant variance) or $k$ (equivalent to having a perfect estimate). As the observed effects become smaller in size, the minimal detectable true effect size increases: i.e., the effect is ‘washed out’ by the single-trial noise, and therefore the true effect has to be larger to be detected.

As the estimation approach for the minimal detectable Hedges’ g (Section A) across the reviewed studies didn’t account for averaging over epochs, the estimated values of the minimal detectable effect size need to be corrected. Let $d_{0}$ be the minimal detectable effect size in case of no within-participant variability (as obtained from the power estimation function, e.g., in R pwr package). If we now assume averaging over $k$ epochs, the adjusted minimal detectable effect size is equal to:

$$d_{adj}=d_{0}\sqrt{1+\frac{1}{r^{2}k}}$$

Since correction for Hedges’ g depends only on the number of degrees of freedom, and applies equally to both $d_{adj}$ and $d_{0}$, the equation for Hedges’ g is very similar:

$$g_{adj} = g_{0}\sqrt{1+\frac{1}{r^{2}k}}$$

The equations for $d_{adj}$ and $g_{adj}$ imply that the minimal detectable effect size becomes larger if only a limited number of epochs per participant is available. However, the extent of this effect depends on the between-within variability ratio $r$, which is not reported in the reviewed studies. Thus, we varied this ratio in simulations discussed below, considering values of 0.1, 0.25, 0.5, 1, and 2. In practice, the ratio can also be estimated from pilot data based on the SD of HER values within and between participants.

We used the estimated or, where available, reported number of epochs (per participant and condition) and adjusted the original minimal detectable values of Hedges’ g (Fig. 7F) accordingly. As shown in Fig. S2A, the effect of trial averaging noticeably affects the resulting distribution only in the case of high within-participant variability ($r=0.1$). Fig. S2B shows how, for a one-sample t-test, the minimal detectable effect size depends on the number of participants and the number of epochs.

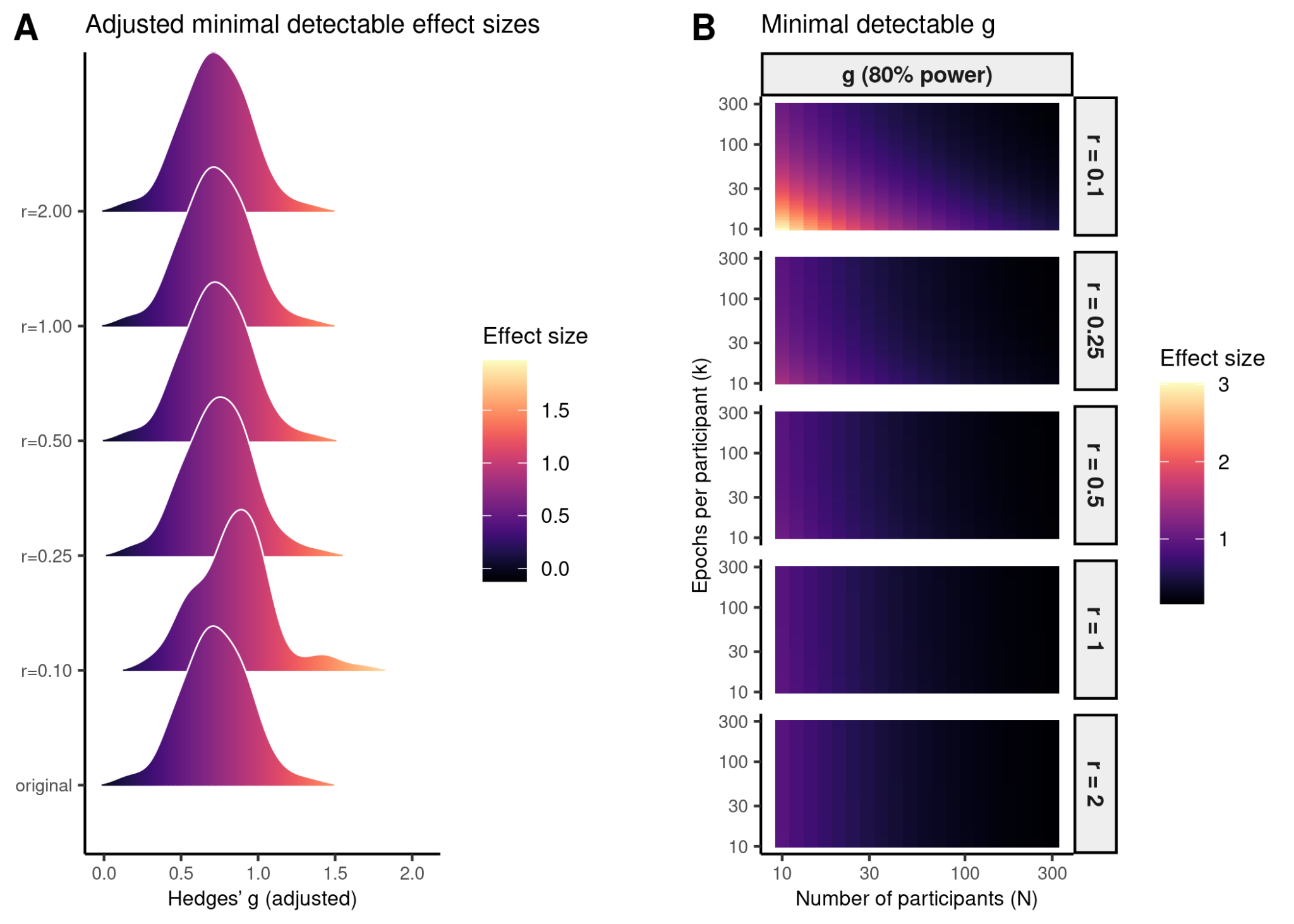

*Figure S2.* Minimal detectable Hedges’ g adjusted for the levels of between- and within-participant variability. (A) The distribution of Hedges’ g that was not adjusted for the number of epochs (similar to Fig. 7F but without correlations), labelled as ‘original’, compared to its adjusted counterparts for different values of between-within variability ratio. (B) The simulated distributions depending on between-within variability ratio, the number of participants, and the number of averaged epochs per participant, for a one-sample t-test. Note that both axes are log-scaled.

### C. A practical note on the required number of trials

The required number of trials depends on the characteristics of recordings, analysis plan and the effects in question (Boudewyn et al., 2018; Gibney et al., 2020). Therefore, one has to consider the specific design to ensure sufficient power, which may pose a challenge. In the following, we describe some examples of how simulations and pilot data can be used to estimate the required number of trials and participants.

#### Simulations

A general approach is to use pilot data, apply all the planned preprocessing steps to it, and resample it many times to create datasets with varying numbers of participants and trials per participant, e.g., with the function sample() in base R. This step is intended to approximate the final data as much as possible and then perform the planned analysis on it to see if the expected effects are detectable and how often.

Sometimes, the pilot data at hand is not an exact replication of the final design (e.g., you plan to compare two conditions but the pilot data contains only one). The goal then is to take neural data with similar characteristics to the planned design and simulate the effect by adding it to the data (like here: Chaumon et al., 2021).

Finally, even if neither neural nor behavioral data are available, we can make some assumptions and simulate datasets (like here: Steinfath et al., 2025). By clearly stating the assumptions and probing the final statistical pipeline, we can already make some approximate conclusions on the power of final analyses. To exemplify, R packages such as fGarch (Wuertz et al., 2023) and MASS (Ripley et al., 2025) can be useful for creating skewed distributions or distributions of multiple correlated variables, but of course other languages such as Matlab and Python provide tools for simulations and resampling as well.

#### Estimating between-within variability from pilot data

The problem with simulation-based approaches is that there is no clear guideline on how to prepare them or assess their quality, and they also may require time-consuming tailoring.

In the previous section B, we propose to use the between-within variability ratio to compute adjusted minimal detectable effect sizes, which is similar to the idea of power contours by Baker et al. (2021). The advantage is that just by knowing the expected within- and between- variabilities of HERs one can compute how exactly the detectable effect sizes will be changing and also assess how adding more participants or trials will affect the power as visualized in Fig. S2B.

As we show in the previous section B, one first has to compute, based on the sample size and a given statistical test, what is the detectable effect size under the no-averaging assumption. This can be done with, for example, the pwr package in R (Champely et al., 2020). Then, one can adjust it based on the between-within ratio.

The between- and within- participant variabilities can be computed using, for example, simple linear mixed-model. Based on the single-trial values per participant, a simple random-intercept model is fitted: e.g., $Signal \sim1+(1|Participant)$. Residual variance is the within-participant estimate, and random-intercept variance is between-participant. Note the method gives $\sigma_{b}^{2}$ and $\sigma_{w}^{2}$. To obtain $r$, one needs to take a square root, or otherwise use directly as $r^{2}$.

For example, Table S1 shows, for a one-sample test and popular experimental settings in the review (number of participants 30 or 60, number of epochs 300), how the different Cohen’s d effect sizes $d_{true}$ in the data (small 0.2, medium 0.5, or large 0.8) translate into the actually observed effect sizes $d_{obs}$ given a medium or small $r^{2}$ of 0.1 or 0.01 ($r$ 0.3 or 0.1), as well as the power to observe these effects. The table also contains example estimates of minimal effect sizes ${mind}_{adj}$ that can be detected with 80% power given the number of participants and epochs.

Note that (1) for small effect sizes, neither 30 nor 60 participants is sufficient for 80% power, and increasing number of trials cannot solve this, for any $r^{2}$; (2) the popular choice of 300 trials (per participant, per condition) is generally sufficient to compensate for trial averaging for $r^{2}$ of 0.1 ($d_{true}$ close to $d_{obs}$), yet in the larger-noise scenario of $r^{2}$ 0.01, the experiments could benefit from more trials; (3) the drastic effects of trial averaging are especially pronounced at lower trial counts, e.g., with 10 trials the actual true effect present in the data $d_{true}$ of 0.8 turns into observed 0.56 or even 0.24 $d_{obs}$ for $r^{2}$ of 0.1 or 0.01, respectively. Finally, even though we do not discuss here the actual between-within variability of HERs, it is already clear that the trial count benchmarks for other popular evoked responses (e.g., P300) are unsuitable for HER studies, given the differences in amplitudes and likely in $r^{2}$.

*Table S1*. Example correspondence between design settings and detectable effect sizes. For one sample test, N - number of participants, d_true - the effect size that we are actually studying, observable under the perfect scenario of an infinite number of epochs, d_obs - the effect that we actually observe after adjusting for the between-within variability and number of epochs, min_d_adj - adjusted minimal detectable effect size given the current settings, including number of epochs, to achieve the power of 0.8.

| N | Epochs | r2 | d_true | d_obs | power | min_d_adj |
| --- | --- | --- | --- | --- | --- | --- |
| 30 | 300 | 0.01 | 0.2 | 0.17 | 0.15 | 0.61 |
| 60 | 300 | 0.01 | 0.2 | 0.17 | 0.26 | 0.42 |
| 30 | 300 | 0.01 | 0.5 | 0.43 | 0.63 | 0.61 |
| 60 | 300 | 0.01 | 0.5 | 0.43 | 0.9 | 0.42 |
| 30 | 300 | 0.01 | 0.8 | 0.69 | 0.95 | 0.61 |
| 60 | 300 | 0.01 | 0.8 | 0.69 | 0.99 | 0.42 |
| 30 | 300 | 0.1 | 0.2 | 0.19 | 0.18 | 0.53 |
| 60 | 300 | 0.1 | 0.2 | 0.19 | 0.32 | 0.37 |
| 30 | 300 | 0.1 | 0.5 | 0.49 | 0.74 | 0.53 |
| 60 | 300 | 0.1 | 0.5 | 0.49 | 0.96 | 0.37 |
| 30 | 300 | 0.1 | 0.8 | 0.78 | 0.98 | 0.53 |
| 60 | 300 | 0.1 | 0.8 | 0.78 | 0.99 | 0.37 |
| 60 | 10 | 0.1 | 0.8 | 0.56 | 0.99 | 0.52 |
| 60 | 10 | 0.01 | 0.8 | 0.24 | 0.45 | 1.2 |

### D. Other supplementary figures and tables

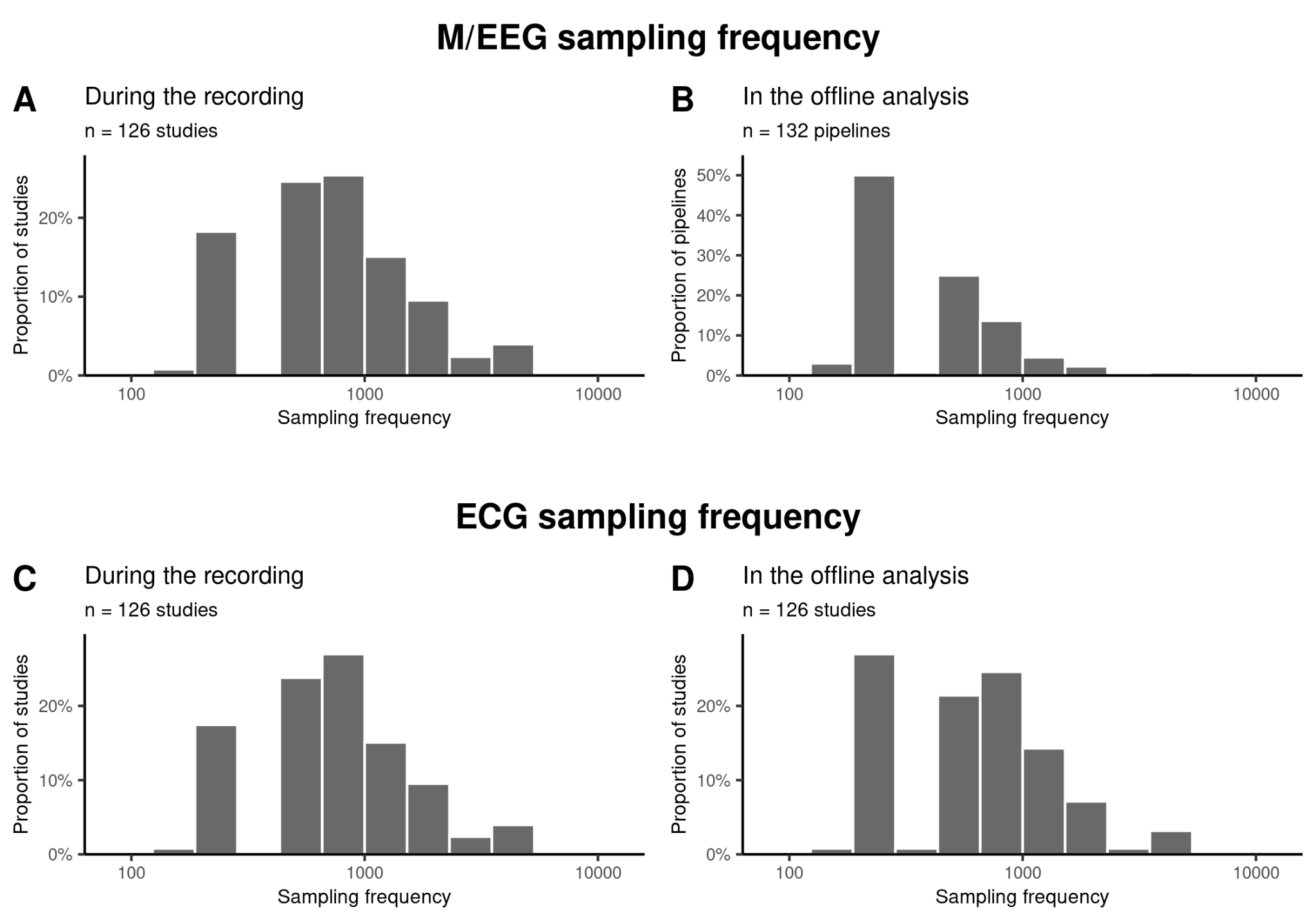

*Figure S3.* Quality control: sampling frequency of M/EEG and ECG was at least 100 Hz in all studies, which reported the respective values. (A) Histogram of the sampling frequency of M/EEG recordings. (B) Histogram of M/EEG sampling frequency in the offline analysis. If no downsampling was mentioned in the reviewed studies, we assumed that it was not performed. Pipelines are shown since one study analyzed two datasets with different sampling frequencies after downsampling. (C-D) Same as A and B but for ECG. If the sampling frequency for ECG was not mentioned explicitly, we assumed that it was the same as for M/EEG.

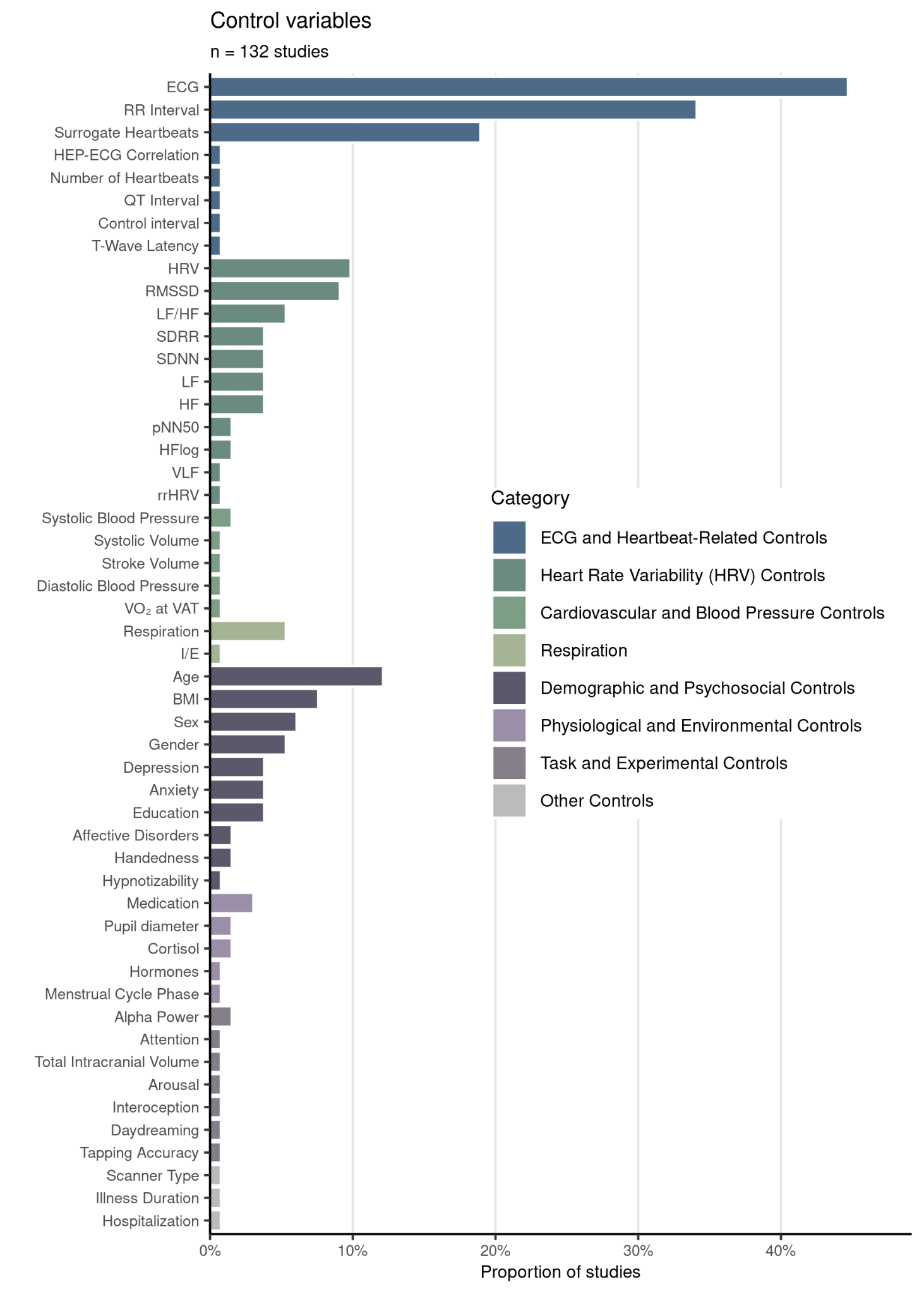

*Figure S4.* Control variables used across pipelines.

Diverse variables are considered to control for spurious findings. The majority reflect heartbeat and cardiovascular system related variables. HR: Heart Rate, SDRR: Standard Deviation of RR intervals, SDNN: Standard deviation of NN intervals, RMSSD: Root Mean Square of Successive Differences, pNN50: Percentage of NN Intervals > 50ms Differences, VLF: Very Low Frequency, LF: Low Frequency, HF: High Frequency, LF/HF: Low Frequency / High Frequency Ratio, HFLog: Log-transformed High Frequency, HRV: Heart Rate Variability, rrHRV: RR Interval-based Heart Rate Variability, VO₂ at VAT: Oxygen Consumption at Ventilatory Anaerobic Threshold, BMI: Body Mass Index, I/E: Inhalation/Exhalation ratio

*Table S2.* List of journals and the number of times they appear among the reviewed studies.

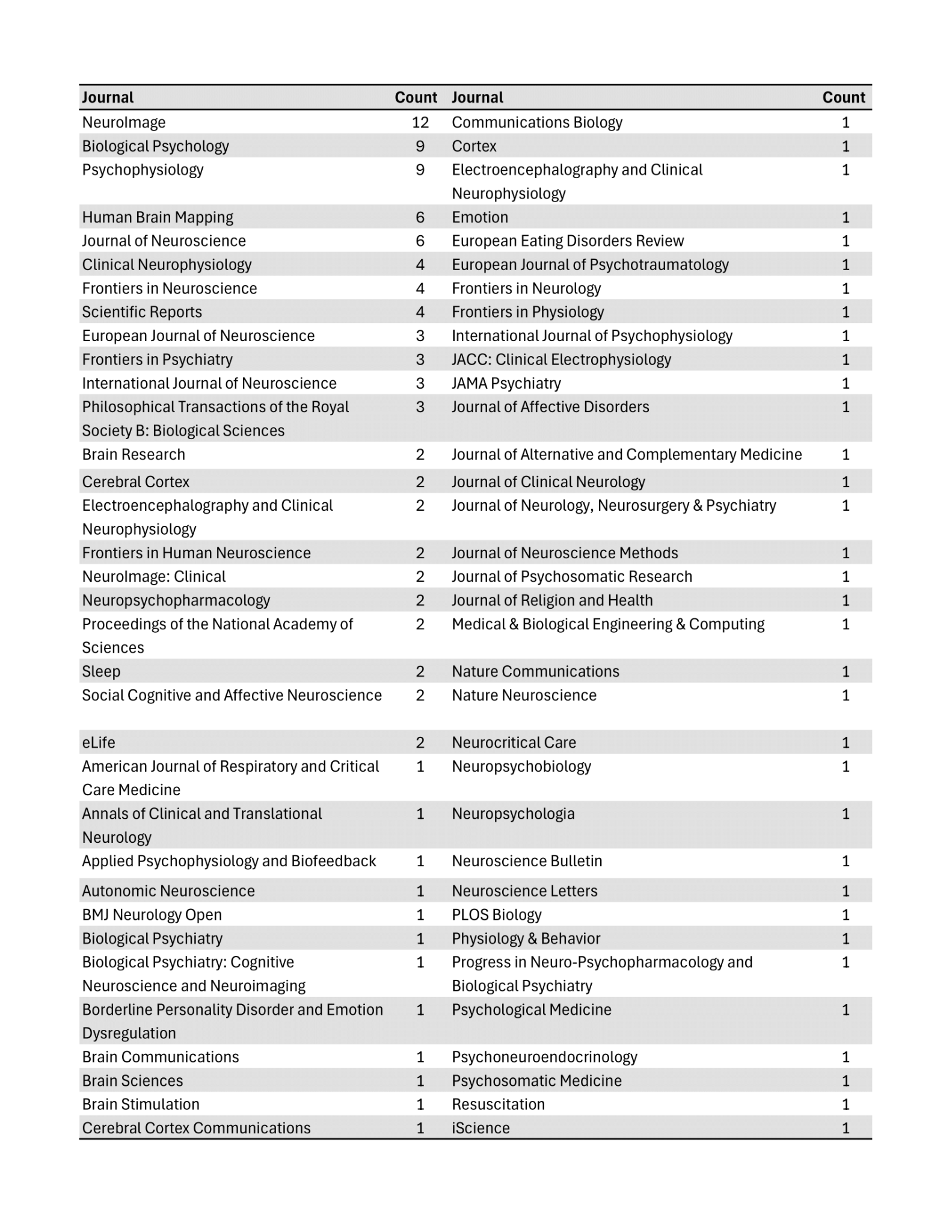

*Table S3*. Overview of the MEG sensors used for data recording and selected for the HER analysis.

| **Paper** | **Layout** | **Grad** | **Mag** | **Sum** | **Selected MEG sensors** |
| --- | --- | --- | --- | --- | --- |
| Zhang et al. (2023) | Elekta | 204 | 102 | 306 | Magnetometers |
| Buot et al. (2021) | Elekta | 204 | 102 | 306 | All |
| Azzalini et al. (2021) | Elekta | 204 | 102 | 306 | Magnetometers |
| Kato et al. (2020) | Elekta | 204 | 102 | 306 | All |
| Babo-Rebelo et al. (2019) | Elekta | 204 | 102 | 306 | Magnetometers |
| Kim et al. (2019) | KRISS | 152 | 0 | 152 | All |
| Babo-Rebelo et al. (2016a) | Elekta | 204 | 102 | 306 | All |
| Babo-Rebelo et al. (2016b) | Elekta | 204 | 102 | 306 | Magnetometers |
| Park et al. (2014) | Elekta | 204 | 102 | 306 | Magnetometers |

### E*.* Details on evaluation of gender composition within HER research

To contextualize gender distribution in HER research within the broader neuroscience field, we subsampled journals according to Dworkin et al. (2020) and Hefter et al. (2025), which had both ranked neuroscience journals by Eigenfactor scores (Bergstrom et al., 2008) from Thomson Reuters’ Web of Science (WoS). Their selection in 2020 included the five highest-ranking neuroscience journals from 1995–2018 (Procedure 1) and in 2025 the 50 highest-ranking ones from 2023 (Procedure 2). Since few of the reviewed HER papers had been published in those journals (Procedure 1: n = 19; Procedure 2: n = 43), we also included the papers from journals with latest Eigenfactor scores (from the WoS’ 2024 Journal Citation Report; Clarivate, 2024) above Hefter et al.’s lowest cutoff (Procedure 3: n = 77). This way we found that authors with female names were more prevalent in higher-ranked journals, decreasing as lower-ranked journals were included (Procedure 1: 84.21% > Procedure 2: 69.77%; Procedure 3: 66.23%; Procedure 3 [Δ remaining papers]: 19.48% [-2.50] FF, 10.39% [-0.96%] MF, 36.6% [-13.72%] FM, 33.77% [+17.18%] MM). Nonetheless, the observed trend more closely resembles the gender distribution (18% FF, 14% MF, 29% FM, 39% MM) that resulted from the large-scale analyses by Hefter et al. (2025).
